# A multidimensional investigation of sleep and biopsychosocial profiles with associated neural signatures

**DOI:** 10.1101/2024.02.15.580583

**Authors:** Aurore A. Perrault, Valeria Kebets, Nicole M. Y. Kuek, Nathan E. Cross, Rackeb Tesfaye, Florence B. Pomares, Jingwei Li, Michael W.L. Chee, Thien Thanh Dang-Vu, B.T. Thomas Yeo

## Abstract

Sleep is essential for optimal functioning and health. Interconnected to multiple biological, psychological and socio-environmental factors (i.e., biopsychosocial factors), the multidimensional nature of sleep is rarely capitalized on in research. Here, we deployed a data-driven approach to identify sleep-biopsychosocial profiles that linked self-reported sleep patterns to inter-individual variability in health, cognition, and lifestyle factors in 770 healthy young adults. We uncovered five profiles, including two profiles reflecting general psychopathology associated with either reports of general poor sleep or an absence of sleep complaints (i.e., sleep resilience) respectively. The three other profiles were driven by the use of sleep aids and social satisfaction, sleep duration and cognitive performance, and sleep disturbance linked to cognition and mental health. Furthermore, identified sleep-biopsychosocial profiles displayed unique patterns of brain network organization. In particular, somatomotor network connectivity alterations were involved in the relationships between sleep and biopsychosocial factors. These profiles can potentially untangle the interplay between individuals’ variability in sleep, health, cognition and lifestyle — equipping research and clinical settings to better support individual’s well-being.

## INTRODUCTION

Decades of research have established that sleep is interconnected to multiple biological, psychological and socio-environmental factors (i.e., biopsychosocial factors)^1–4^. Importantly, sleep difficulties are among the most common comorbidities of mental and physical disorders^5–8^, highlighting the central role of sleep in health. Despite the recognition that sleep is a unique marker for optimal health^9,10^ and a potential transdiagnostic therapeutic target, its multidimensional and transdisciplinary nature is rarely capitalized on in research. Traditionally, single-association studies have investigated the relationship between a single dimension of sleep (e.g., duration, quality, onset latency) and/or a single outcome of interest. Such uni-dimensional studies have demonstrated links between insufficient or poor sleep with a multitude of negative outcomes separately, including cognitive difficulties^11,12^, brain connectivity changes^13–15^, decreased physical health^7,16^, mental health and well-being^8,17^, as well as increased risks for cardiovascular disease^7,18,19^, neurodegenerative disease^20,21^ and psychiatric disorders^8,22^. However, by treating sleep as a binary domain (e.g., good vs. poor sleep, short vs. long), these studies fail to capture the multidimensional nature of sleep and the multiple intricate links with biological, psychological, and socio-environmental (i.e., biopsychosocial) factors. Therefore, it remains unclear which biopsychosocial factors are most robustly associated with sleep traits and whether these factors are supported by similar neural mechanisms.

Adding to the complexity of these relationships is how sleep and good sleep health are defined. Characterizing sleep is a challenging task because of its multidimensional nature^23^. Sleep can be defined by its quantity (i.e., sleep duration) and quality (i.e., satisfaction, efficiency), as well as in terms of regularity, timing, and alertness. These dimensions are deemed particularly relevant when defining sleep health^9^, as they each have been related to biopsychosocial outcomes. Different sleep dimensions can also be described as either “good” or “bad” sleep, without necessarily affecting one another, e.g., short sleep duration is not systematically associated with poor sleep quality. Another important aspect of sleep is how it is subjectively characterized. For instance, our perception of sleep can influence daytime functioning^24^ and can be ascribed to certain behaviors that differ from objective reports^25,26^. Reconciling the multiple components of sleep and the complex connections to a myriad of biopsychosocial factors requires frameworks grounded in a multidimensional approach. The biopsychosocial model has long been used to assert that biological (e.g., genetics and intermediate brain phenotypes), psychological (e.g., mood and behaviors), and social factors (e.g., social relationships, economic status), are all significant contributors to health and disease^2,3^. Indeed, the biopsychosocial model has been used to establish current diagnostic and clinical guidelines, such as the World Health Organization’s International Classification of Functioning, Disability and Health, and is considered central to person-centered care^27^. Hence, statistical methods that enable us to interrogate the complex interconnected relationships within and between sleep and biopsychosocial factors can advance our understanding of optimal health and functioning across the lifespan. Multivariate data-driven techniques can help disentangle these complex interrelations, by deriving latent components that optimally relate multidimensional data sets in a single integrated analysis. A few studies have used such techniques to account for the multidimensional components of sleep and biopsychosocial factors separately^15,28–32^. However, no study has integrated both multidimensional components of sleep and biopsychosocial factors to derive profiles that can account for the dynamic interplay among biopsychosocial factors, and link such components with brain network organization.

Deploying multivariate data-driven techniques requires a large sample size to identify latent components that can be generalised well^33–35^. One such optimal dataset is the Human Connectome Project dataset (HCP)^36^ as it comprises a wide range of self-reported questionnaires about lifestyle, mental and physical health, personality and affect, as well as objective measures of physical health and cognition from over a thousand healthy young adults. Moreover, the HCP dataset stands out as one of the rare large-scale datasets that implemented a detailed assessment of sleep health, i.e., the Pittsburg Sleep Quality Index (PSQI)^37^. This standardized sleep questionnaire, used both by clinicians and researchers, assesses different dimensions of sleep health in 19 individual items, creating 7 sub-components defining different dimensions of sleep, including sleep duration, satisfaction, and disturbances.

Beyond sleep-biopsychosocial profiling, the HCP dataset also provides the opportunity to explore the neural signatures of these sleep-biopsychosocial profiles using magnetic resonance imaging (MRI). Multiple studies have shown that neural signal fluctuation patterns during rest (i.e., resting-state functional connectivity; RSFC) are sensitive to sleep dimensions (e.g., sleep duration, sleep quality) ^14,15,32,38^, but also predictive of psychopathology (e.g., depressive symptoms, impulsivity)^39,40^ and cognitive performance^14,38^. However, the way large-scale network organization may differentially affect individuals’ variability in sleep, psychopathology, cognition and lifestyle, remains to be characterized beyond unidimensional association studies. Such holistic biopsychosocial approaches are not only in line with established diagnostic frameworks but also with initiatives such as the NIMH’s Research Domain Criteria (RDoC) that encourage investigating mental disorders as continuous dimensions rather than distinct categories by integrating data from genomics, neural circuitry and behavior^41–43^.

Identifying vulnerability markers constitutes a first step towards forecasting disease trajectories and designing multimodal multidimensional targeted therapies. Given the increasing recognition that sleep has a central role in health and well-being, we believe that sleep profiles should be included as a core aspect of these markers. Hence, in this study, we sought to take a multidimensional data-driven approach to identify sleep-biopsychosocial profiles that simultaneously relate self-reported sleep patterns to biopsychosocial factors of health, cognition, and lifestyle in the HCP cohort of healthy young adults^36^. We further explored patterns of brain network organization associated with each profile to better understand their neurobiological underpinnings.

## RESULTS

We applied canonical correlation analysis (CCA) to derive latent components (LCs) linking the 7 sub-components of the PSQI to 118 biopsychosocial measures (spanning cognitive performance, physical and mental health, personality traits, affects, substance use, and demographics; Table S1) in 770 healthy adults from the S1200 release of the HCP dataset^36^ (Figure 1A). Participants were young adults between 22 and 36 years old (mean 28.86 ± 3.61 years old, 53.76% female), were generally employed full-time (70.7%) and were mostly white (78%; see Table 1 for Demographics).

**Figure 1.**
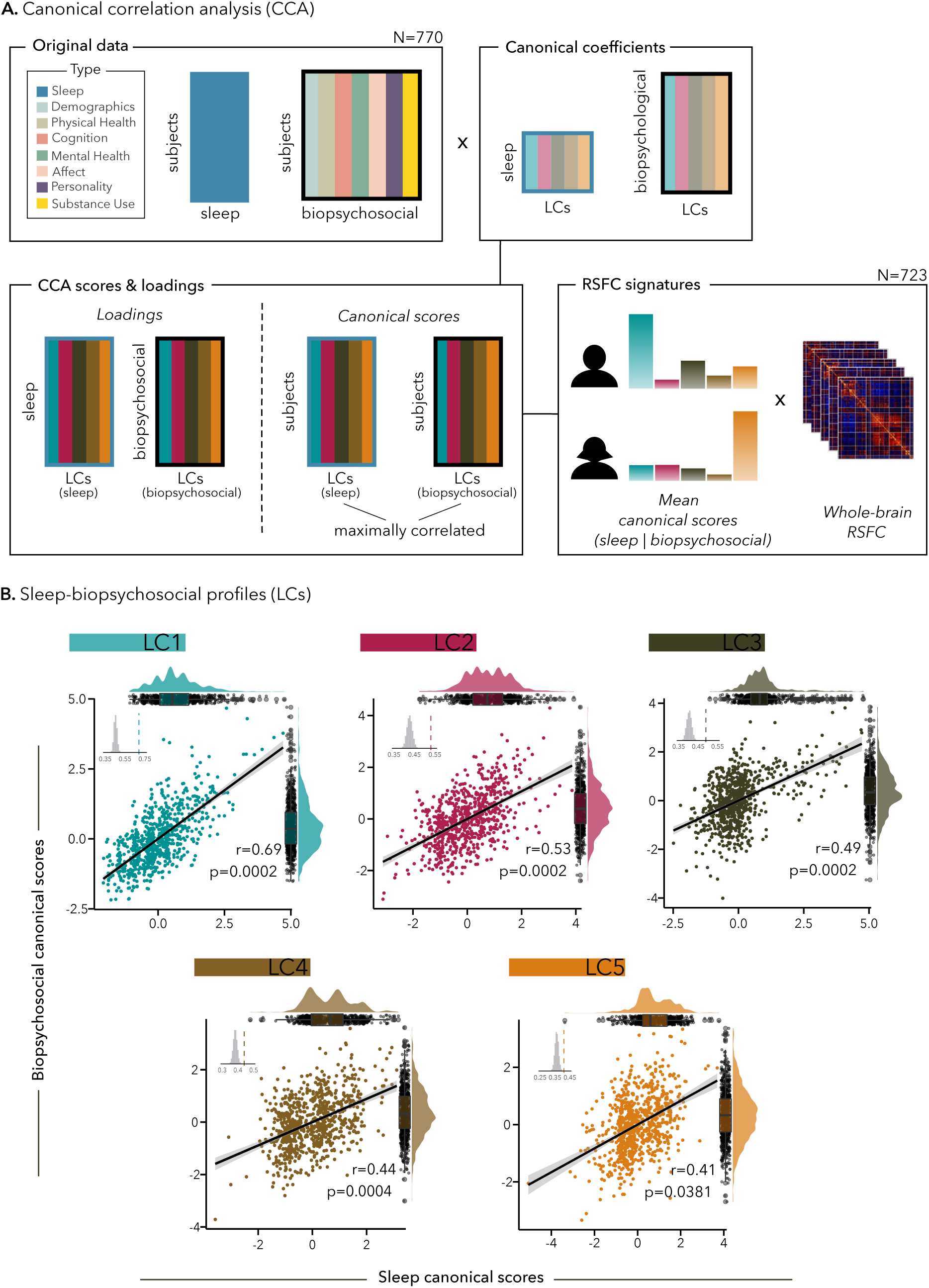
Canonical correlation analysis reveals five sleep-biopsychosocial profiles (LCs). (A) Canonical correlation analysis (CCA) flowchart and RSFC signatures; (B) Scatter plots showing correlations between biopsychosocial and sleep canonical scores. Each dot represents a different participant. Inset shows the null distribution of canonical correlations obtained by permutation testing; note that the null distribution is not centered at zero. The dashed line indicates the actual canonical correlation computed for each LC. The distribution of sleep (top) and biopsychosocial (right) canonical scores is shown on rain cloud plots.

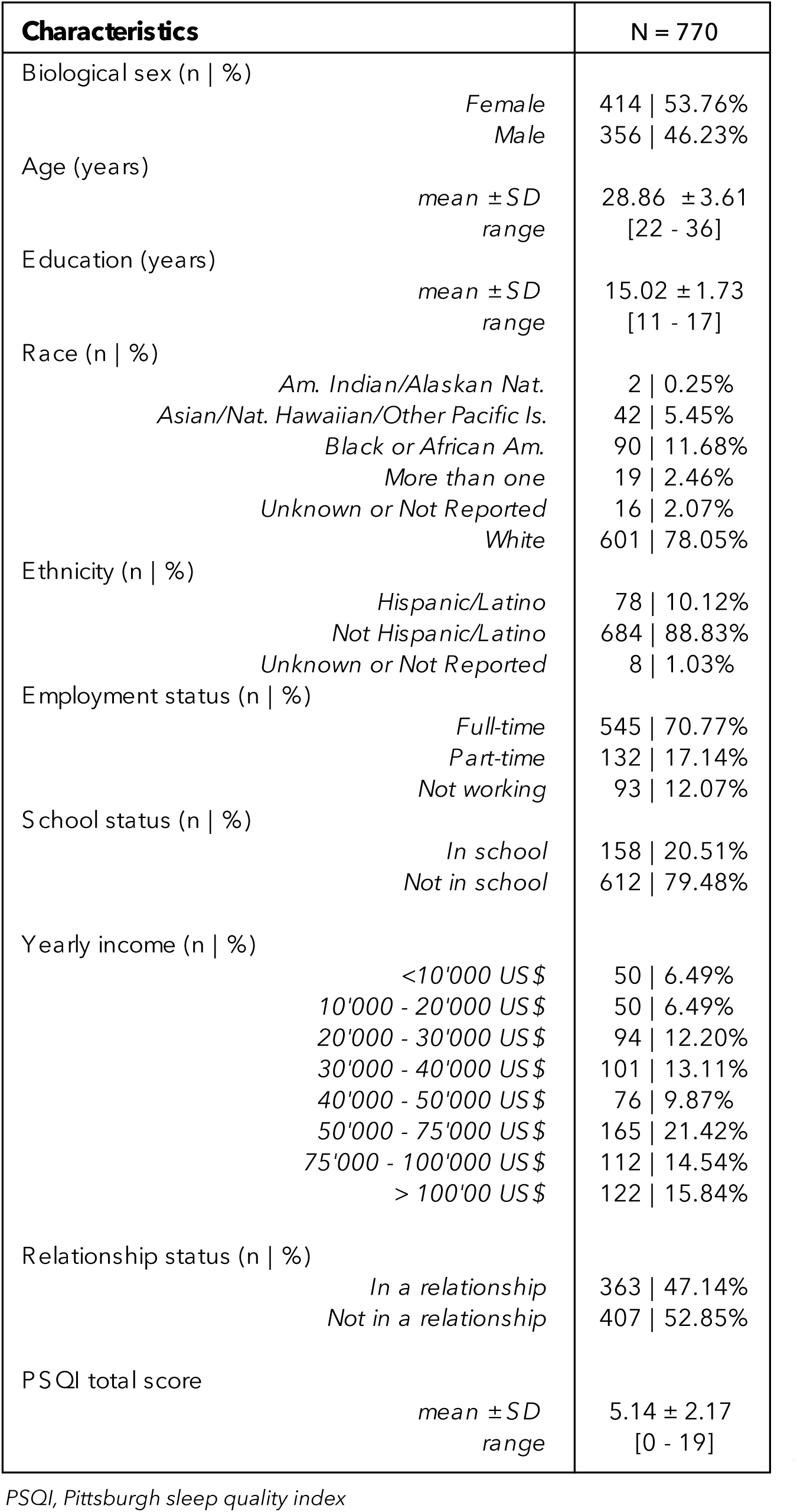

### Five latent components (LCs) linking sleep and biopsychosocial factors

Out of the seven significant LCs that were derived, 5 were interpretable LCs delineating multivariate relationships between sleep and biopsychosocial factors (Figure 1B; a description of LC6 and LC7 can be found in the Supplementary Material – Figure S1). While LC1 and LC2 were defined by general patterns of sleep (either general poor sleep or sleep resilience), LCs 3-5 reflected more specific sub-components of the PSQI, all associated with specific patterns of biopsychosocial factors. The 5 LCs respectively explained 88%, 4%, 3%, 2%, 1% of covariance between the sleep and biopsychosocial data.

LC1 was characterized by a general pattern of poor sleep, including decreased sleep satisfaction, longer time to fall asleep, greater complaints of sleep disturbances and daytime impairment, as well as greater (i.e., worse) psychopathology (e.g., depression, anxiety, somatic complaints, internalizing behavior) and negative affect (e.g., fear, anger, stress – Figure 2A).

**Figure 2.**
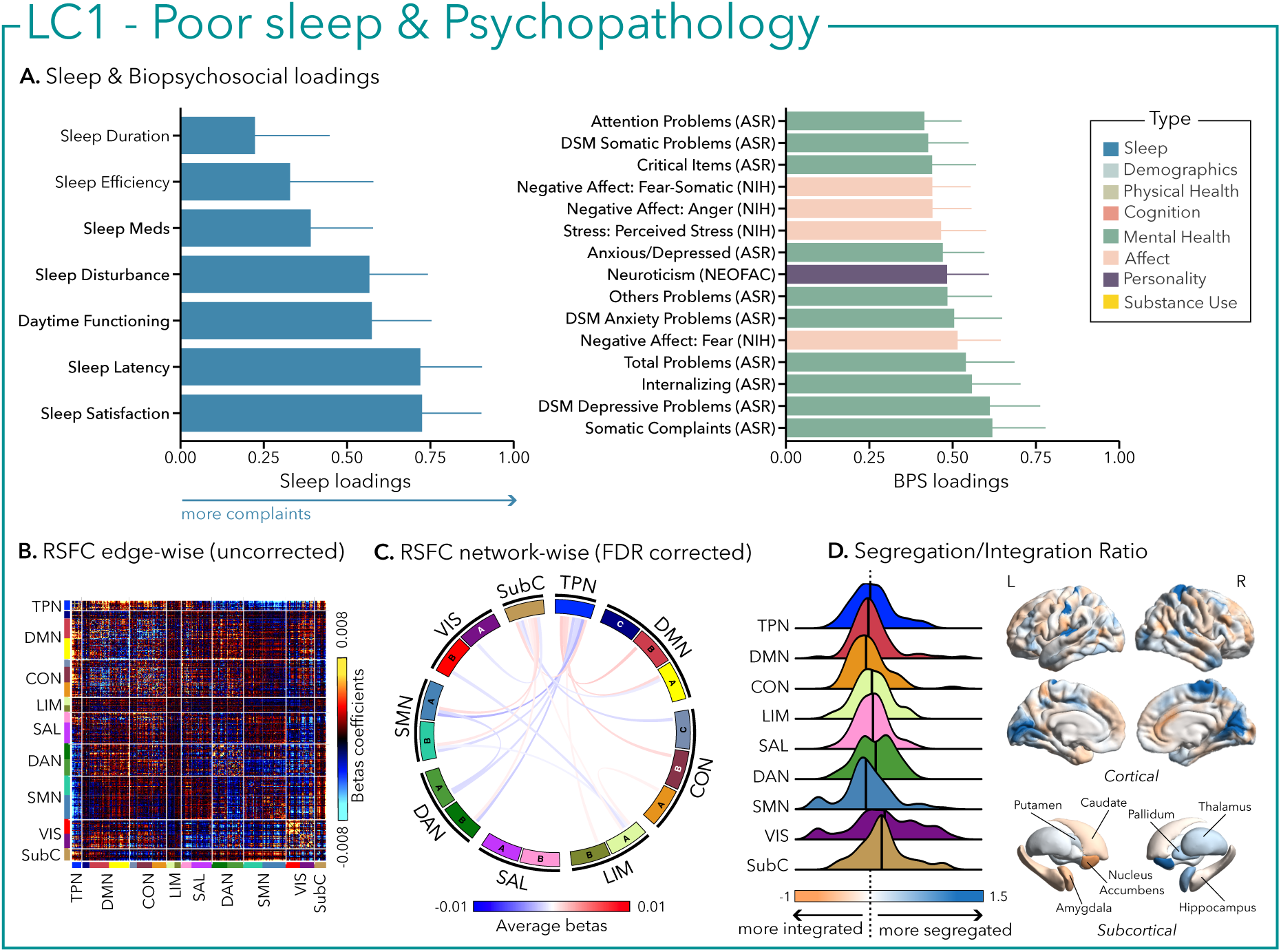
The first latent component (LC1) reflects poor sleep and psychopathology. (A) Sleep loadings (left) and top 15 strongest biopsychosocial (BPS) loadings (right) for LC1. Greater loadings on LC1 were associated with higher measures of poor sleep and psychopathology. Higher values on sleep (blue) and biopsychosocial (green, purple, pink) loadings indicate worse outcomes. Error bars indicate bootstrapped-estimated confidence intervals (i.e., standard deviation) and measures in bold indicate statistical significance (after FDR correction q<0.05); (B) Unthresholded edge-wise beta coefficients obtained from generalized linear models (GLM) between participants’ LC1 canonical scores (i.e., averaged sleep and biopsychosocial canonical scores) and their RSFC data; (C) FDR-corrected network-wise beta coefficients computed with GLMs within and between 17 large-scale brain networks^44^ and subcortical regions^45^. (D) Distribution of the integration/segregation ratio in each of the 7 large-scale brain networks and subcortical regions associated with LC1 (left). The dashed line indicates the median of all parcels, and the bold black lines represent the median for each network. The integration/segregation ratio values for the 400 Schaeffer parcellation^46^ and 7 subcortical regions are projected on cortical and subcortical surfaces (right).

Similarly, LC2 was also driven by greater psychopathology, especially attentional problems (e.g., inattention, ADHD), low conscientiousness, and negative affect (Figure 3A). In terms of sleep, however, in contrast to the first LC, greater psychopathology was only related to higher complaints of daytime impairment without complaints of sleep difficulties, suggesting sleep resilience.

**Figure 3.**
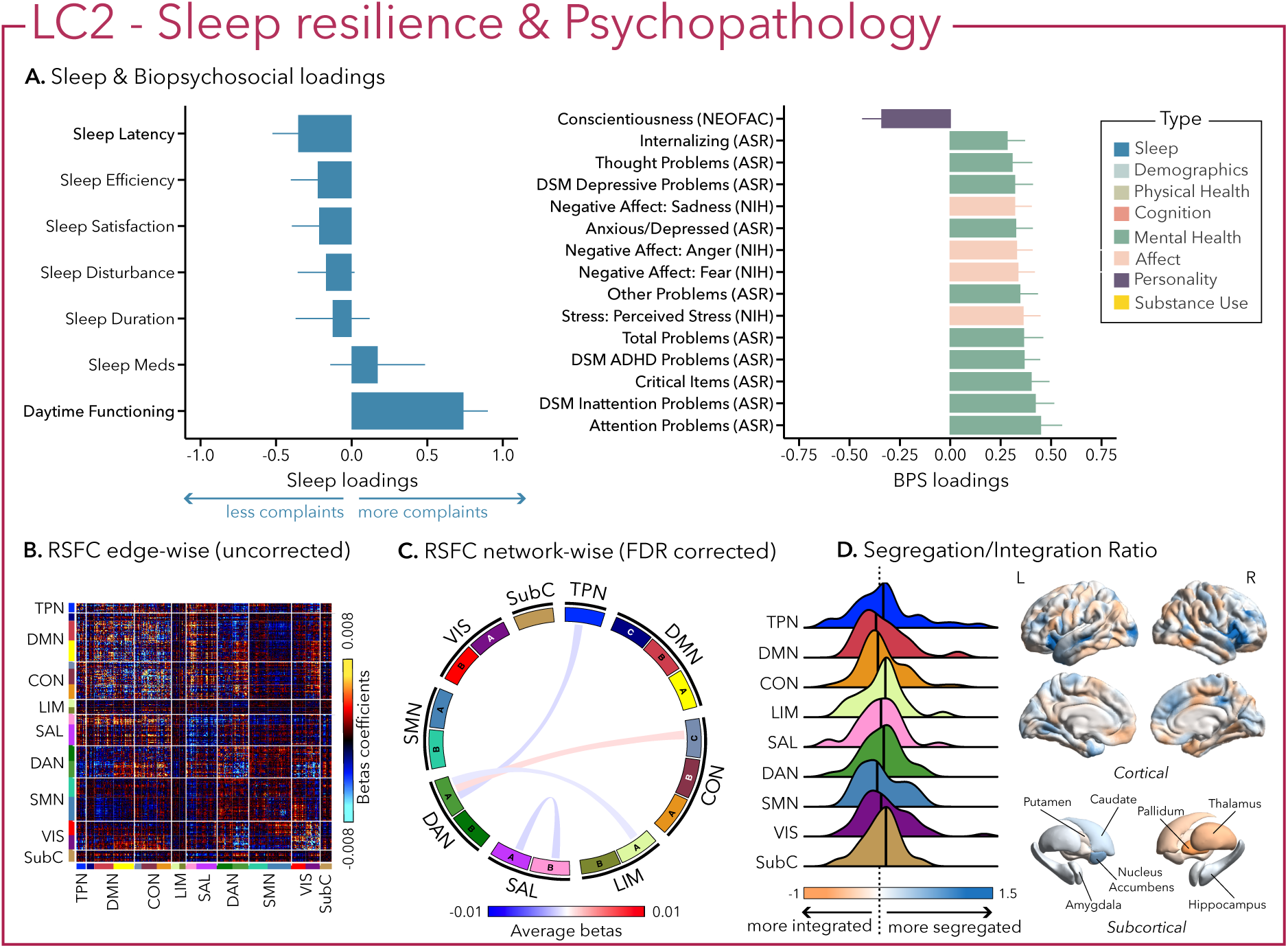
The second latent component (LC2) reflects sleep resilience and psychopathology. (A) Sleep loadings (left) and top 15 strongest biopsychosocial (BPS) loadings (right) for LC2. Greater loadings on LC2 were associated with higher measures of complaints of daytime dysfunction and psychopathology. Positive values on sleep (blue) loadings indicate worse outcomes while positive values on biopsychosocial (green, purple, pink) loadings reflect higher magnitude on these measures. Error bars indicate bootstrapped-estimated confidence intervals (i.e., standard deviation) and measures in bold indicate statistical significance. (B) Unthresholded edge-wise beta coefficients obtained from generalized linear models (GLM) between participants’ LC2 canonical scores (i.e., averaged sleep and biopsychosocial canonical scores) and their RSFC data; (C) FDR-corrected network-wise beta coefficients computed with GLMs within and between 17 large-scale brain networks^44^ and subcortical regions^45^. (D) Distribution of the integration/segregation ratio in each of the 7 large-scale brain networks and subcortical regions associated with LC2 (left). The dashed line indicates the median of all parcels, and the bold black lines represent the median for each network. The integration/segregation ratio values for the 400 Schaeffer parcellation^46^ and 7 subcortical regions are projected on cortical and subcortical surfaces (right).

LC3 was mostly characterized by sleep-aids intake (i.e., sleep meds PSQI sub-component) and to a lesser extent a lack of daytime functioning complaint. Surprisingly, LC3 was not driven by any attentional problem but was related to worse performance in visual episodic memory and emotional recognition. Moreover, sleep aids/hypnotics intake was mainly related to satisfaction in social relationships (Figure 4A).

**Figure 4.**
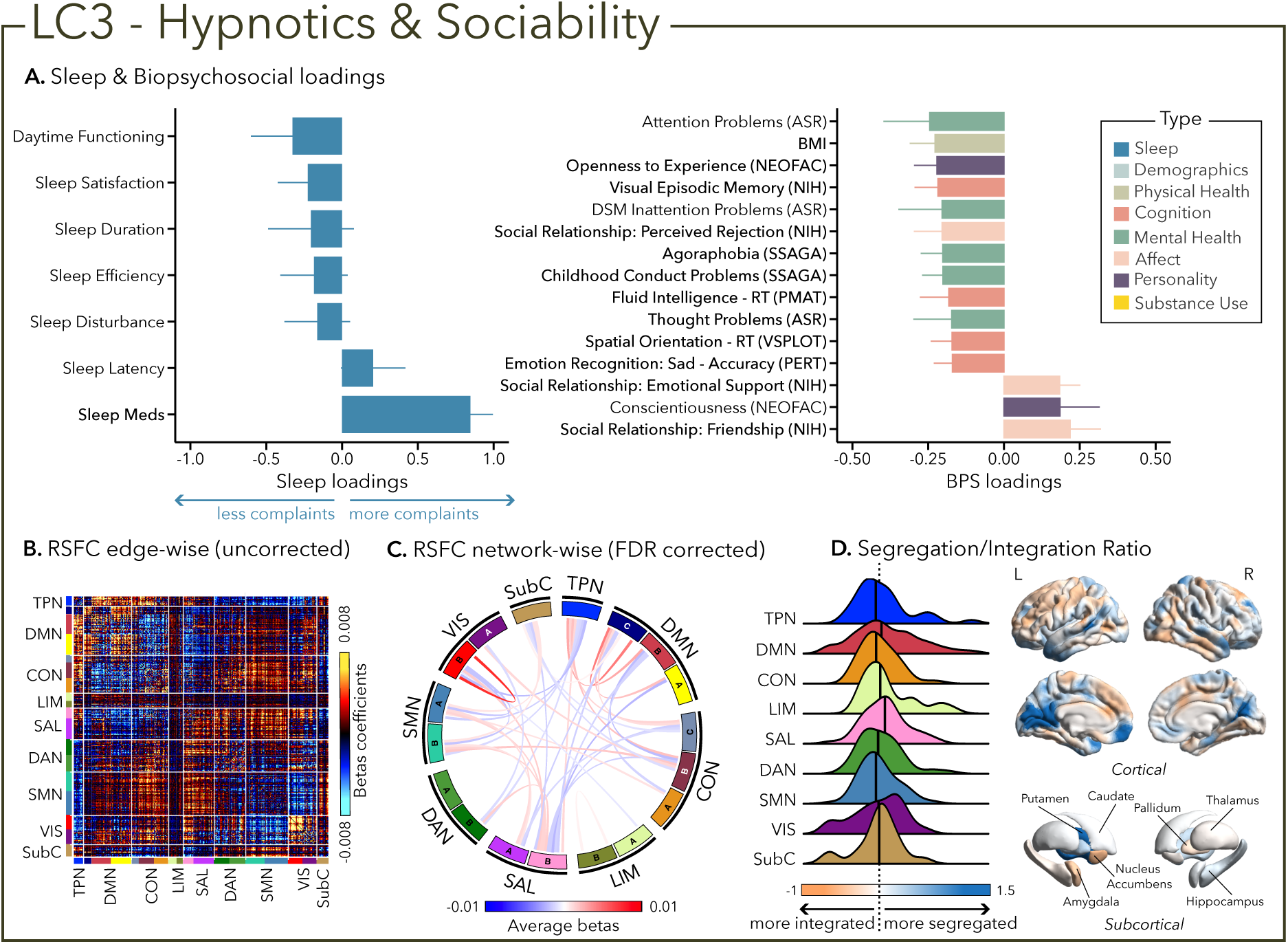
The third latent component (LC3) reflects hypnotics and sociability. (A) Sleep loadings (left) and top 15 strongest biopsychosocial (BPS) loadings (right) for LC3. Greater loadings on LC3 were associated with the use of sleep aids and measures of positive social relationships, lower body mass index (BMI) and poor visual episodic memory performance. Positive values on sleep (blue) loadings indicate worse outcomes while positive values on the mental health (green), affect (pink) and personality (purple) categories of biopsychosocial loadings reflect higher magnitude on these measures. Positive value in the physical health (olive) category represents higher value and positive values in the cognition (orange) category indicate either higher accuracies or slower reaction times (RT). Error bars indicate bootstrapped-estimated confidence intervals (i.e., standard deviation) and measures in bold indicate statistical significance. (B) Unthresholded edge-wise beta coefficients obtained from generalized linear models (GLM) between participants’ LC3 canonical scores (i.e., averaged sleep and biopsychosocial canonical scores) and their RSFC data; (C) FDR-corrected network-wise beta coefficients computed with GLMs within and between 17 large-scale brain networks^44^ and subcortical regions^45^. (D) Distribution of the integration/segregation ratio in each of the 7 large-scale brain networks and subcortical regions associated with LC3 (left). The dashed line indicates the median of all parcels, and the bold black lines represent the median for each network. The integration/segregation ratio values for the 400 Schaeffer parcellation^46^ and 7 subcortical regions are projected on cortical and subcortical surfaces (right).

While LC4 was solely driven by sleep duration (i.e., not sleeping enough - reporting <6-7h per night), LC5 was mostly characterized by the presence of sleep disturbances that can encompass multiple awakenings, nocturia and breathing issues as well as pain or temperature imbalance. In LC4, short sleep duration was associated with worse accuracy and longer reaction time at multiple cognitive tasks tapping into emotional processing, delayed reward discounting, language, fluid intelligence, and social cognition. LC4 was also characterized by higher aggressive behavior and lower agreeableness (Figure 5A).

**Figure 5.**
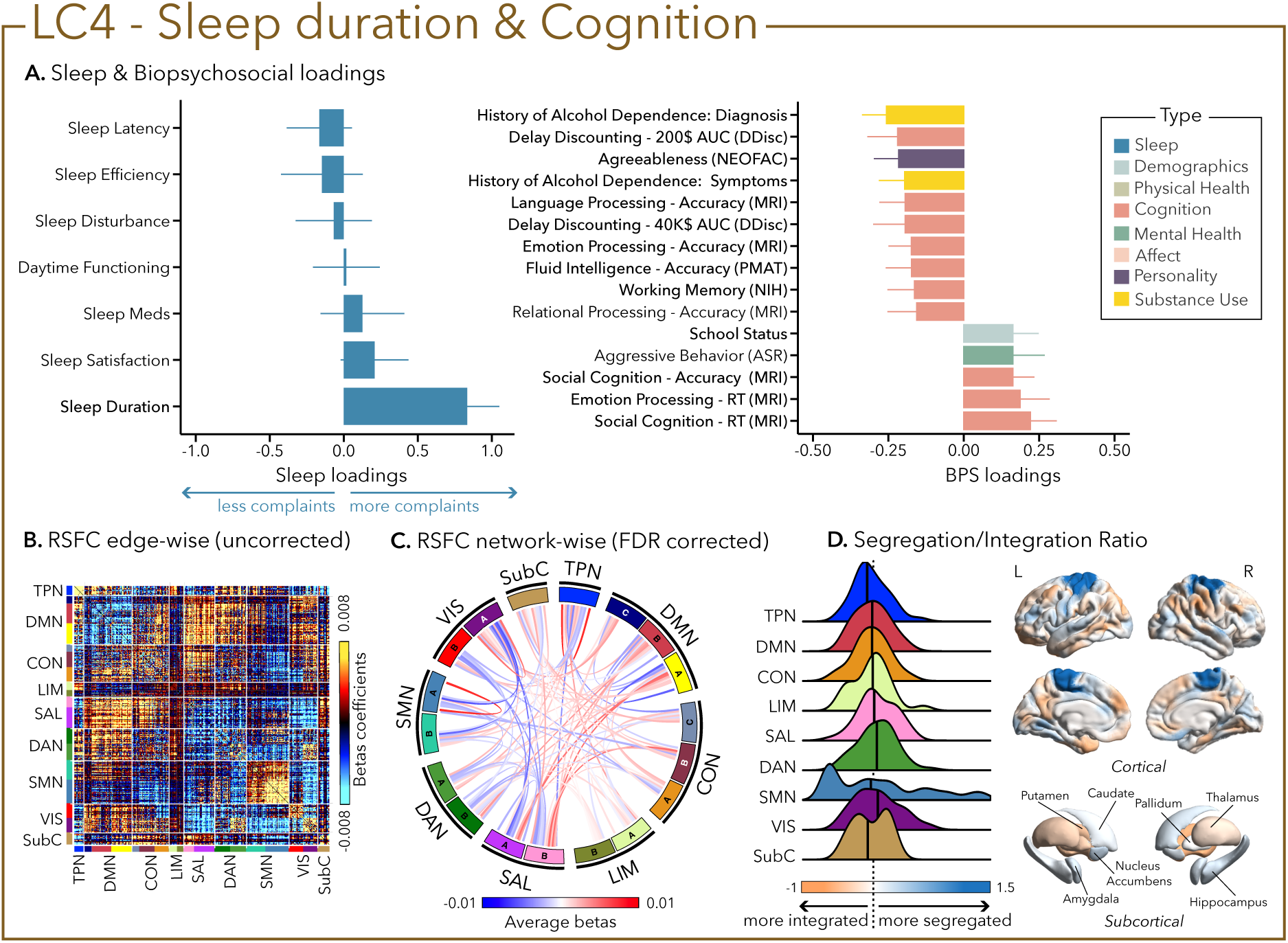
The fourth latent component (LC4) reflects sleep duration and cognition. (A) Sleep loadings (left) and top 15 strongest biopsychosocial (BPS) loadings (right) for LC4. Greater loadings on LC4 were associated with shorter sleep duration and measures of poor cognitive performance. Positive values on sleep loadings indicate worse outcomes while positive values on the mental health (green), substance use (yellow), demographics (light blue) and personality (purple) categories of biopsychosocial loadings reflect higher magnitude on the measures. Positive values in the cognition (orange) category indicate either higher accuracies or slower reaction times (RT). Error bars indicate bootstrapped-estimated confidence intervals (i.e., standard deviation) and measures in bold indicate statistical significance. (B) Unthresholded edge-wise beta coefficients obtained from generalized linear models (GLM) between participants’ LC4 canonical scores (i.e., averaged sleep and biopsychosocial canonical scores) and their RSFC data; (C) FDR-corrected network-wise beta coefficients computed with GLMs within and between 17 large-scale brain networks^44^ and subcortical regions^45^. (D) Distribution of the integration/segregation ratio in each of the 7 large-scale brain networks and subcortical regions associated with LC4 (left). The dashed line indicates the median of all parcels, and the bold black lines represent the median for each network. The integration/segregation ratio values for the 400 Schaeffer parcellation^46^ and 7 subcortical regions are projected on cortical and subcortical surfaces (right).

Interestingly, sleep disturbances in LC5 were also associated with aggressive behavior and worse cognitive performance (e.g., in language processing and working memory), but were mostly characterized by critical items on mental health assessments (i.e., anxiety, thought problems, internalization) and substance abuse (i.e., alcohol and cigarette use – Figure 6A).

**Figure 6.**
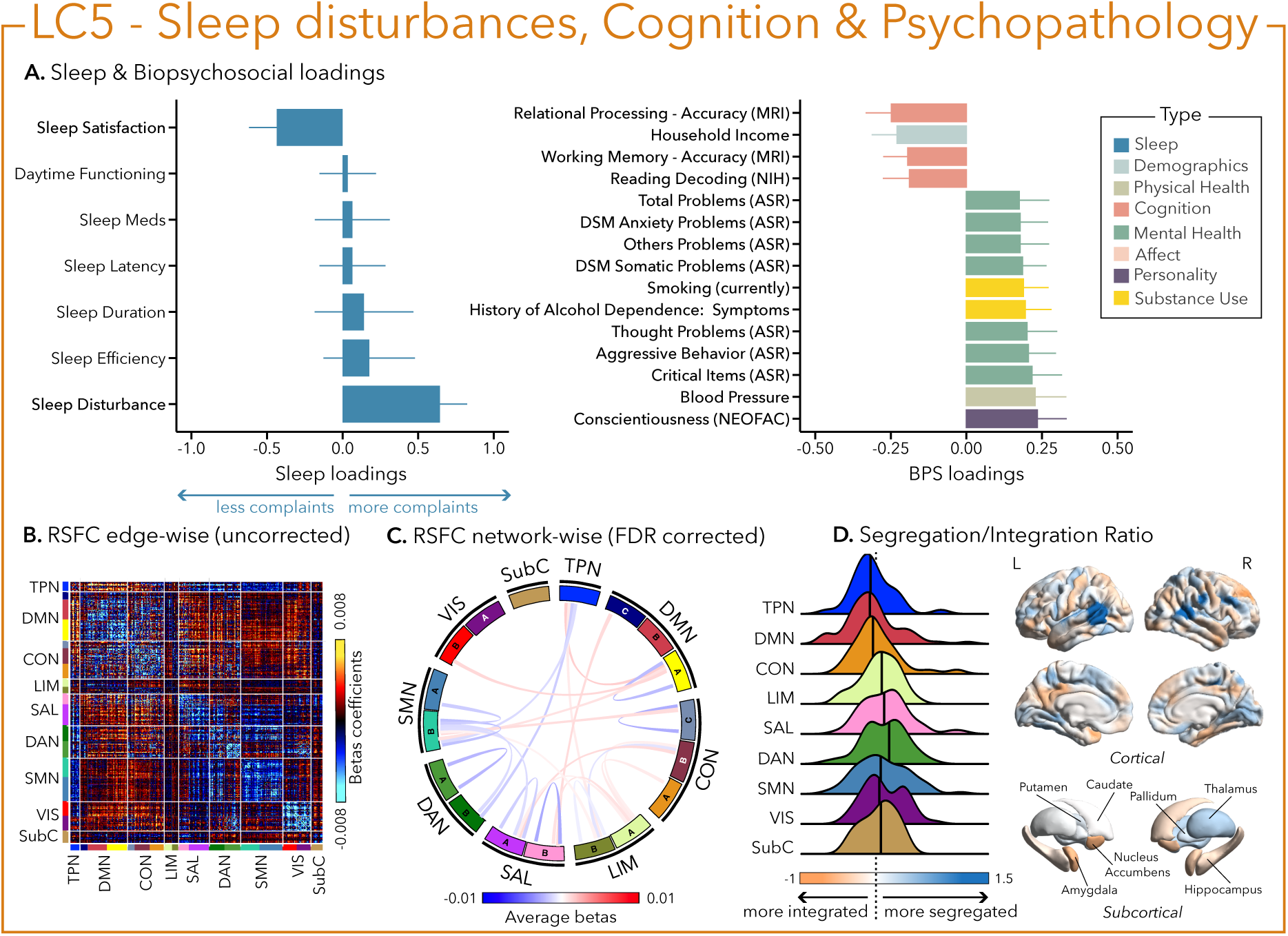
The fifth latent component (LC5) reflects sleep disturbance, cognition and psychopathology. (A) Sleep loadings (left) and top 15 strongest biopsychosocial (BPS) loadings (right) for LC5. Greater loadings on LC5 were associated with the presence of sleep disturbances, higher measures of psychopathology and lower cognitive performance. Positive values on sleep loadings indicate worse outcomes while positive values on the mental health (green), substance use (yellow) and personality (purple) categories of biopsychosocial loadings reflect higher magnitude on these measures. Positive values in the cognition (orange) category indicate either higher accuracies or slower reaction times (RT), while positive values in the demographics (light blue) and physical health (olive) categories represent higher values. Error bars indicate bootstrapped-estimated confidence intervals (i.e., standard deviation) and measures in bold indicate statistical significance. (B) Unthresholded edge-wise beta coefficients obtained from generalized linear models (GLM) between participants’ LC5 canonical scores (i.e., averaged sleep and biopsychosocial canonical scores) and their RSFC data; (C) FDR-corrected network-wise beta coefficients computed with GLMs within and between 17 large-scale brain networks^44^ and subcortical regions^45^. (D) Distribution of the integration/segregation ratio in each of the 7 large-scale brain networks and subcortical regions associated with LC5 (left). The dashed line indicates the median of all parcels, and the bold black lines represent the median for each network. The integration/segregation ratio values for the 400 Schaeffer parcellation^46^ and 7 subcortical regions are projected on cortical and subcortical surfaces (right).

### Sleep and biopsychosocial profiles exhibit distinct signatures of resting-state brain connectivity

In terms of brain organization, the 5 LCs revealed distinct patterns of network connectivity. Specifically, we examined patterns of both within-network and between-network connectivity (see Figure S2 for subcortical-cortical patterns).

Greater (averaged) biopsychosocial and sleep composite scores on LC1 were associated with increased RSFC between subcortical areas and the somatomotor and dorsal attention networks (Figures 2B and 2C), and a decreased RSFC between the temporoparietal network and these two networks. The visual network showed a flattened distribution of segregation/integration ratio (i.e., more variability in segregation and integration among the parcels of the network). The amygdala and nucleus accumbens exhibited asymmetrical patterns in the segregation/integration ratio with the left side being more segregated (Figure 2D). Meanwhile, LC2 was associated with increased RSFC between the dorsal attention and control network but decreased RSFC between dorsal attention and the temporoparietal and limbic networks (Figures 3B and 3C), a higher segregation of nodes within the temporoparietal network and increased integration within the right thalamus (Figure 3D). Higher composite scores in LC3 were associated with increased RSFC within the visual and default mode networks (Figures 4B and 4C). The segregation/integration ratio within the default mode exhibited a flattened distribution (i.e., high variability in segregation and integration among parcels) but there was an increased segregation in the limbic and visual networks (Figure 4D). While greater composite scores in LC4 were associated with widespread patterns of hypo- or hyper-connectivity within and between every network the somatomotor network specifically exhibited an altered pattern of segregation and integration (Figures 5B to 5D). Finally, we found that greater averaged composite scores in LC5 were mainly associated with reduced within-network connectivity in the somatomotor, dorsal and ventral attention networks (Figures 6B and 6C) but no strong pattern of segregation/integration ratio change (Figure 6D).

### Post-hoc associations with socio-demographics, health, and family history of mental health

We found a number of significant associations between LC composite scores and socio-economic (e.g., education level, household income) and socio-demographic factors (e.g., race, ethnicity; see Table S4 and Supplemental Results). In brief, most profiles (LCs 1,4,5) showed significant associations between sleep-biopsychosocial composite scores and education level, where lower education level was associated with a higher composite score in LCs 1,4,5 (all q<0.05). Similarly, lower household income correlated with a higher composite score in LCs 1-2 (all q<0.05). Race and ethnicity groups revealed differences in composite sleep and biopsychosocial scores for LCs 1,3-5 (all q<0.05). Finally, while the presence of a family history of psychopathology was associated with higher biopsychosocial scores in LCs 1-2, we only found biological sex differences in LC5, with higher sleep and biopsychosocial composite scores in female participants (q<0.05).

### Control analyses

We summarize several analyses that demonstrate the robustness of our findings. First, LC1 and LC2 successfully generalized in our cross-correlation scheme (mean across 5 folds: r=0.49, p=0.001; r=0.19, p=0.039 respectively), but not LCs 3-5 (see Table S3), suggesting that LCs 3-5 might not be as robust and generalizable, possibly due to these LCs being driven by a single sleep dimension. Second, we re-computed the CCA analysis after: (i) applying quantile normalization on sleep and biopsychosocial measures; (ii) excluding participants that had tested positive for any substance on the day of the MRI; (iii) excluding physical health measures (i.e., body mass index, hematocrit, blood pressure) or (iv) sociodemographic variables (i.e., employment status, household income, school status, relationship status) from the biopsychosocial matrix. The CCA loadings remained mostly unchanged (Table S5). We also assessed the robustness of our imaging results in several ways. First, we re-computed the GLM analysis using RSFC data that underwent CompCor^47^ instead of GSR. The RSFC patterns were not much altered, as shown by generally high correlations with the main analysis (r=0.75, r=0.76, r=0.78, r=0.51, r=0.77 for LCs 1-5 respectively; Figure S3). Next, excluding subjects that likely fell asleep in the scanner did not impact our findings (r=0.90, r=0.87, r=0.95, r=0.95, r=0.95 for LCs 1-5 respectively; Figure S3); however, we found that these participants had higher sleep and biopsychosocial composite scores on LC4 compared to participants that likely stayed awake during the scan (Figure S4). Finally, we re-computed the GLM analyses by using sleep and biopsychosocial canonical scores instead of averaged scores. We found moderate to high correlations with the main GLM analysis (r=0.69, r=0.62, r=0.63, r=0.46, r=0.67 for LCs 1-5 respectively; Figure S3).

## DISCUSSION

Leveraging a multidimensional data-driven approach in a large cohort of healthy young adults, we uncovered five distinct sleep profiles linked to biopsychosocial factors encompassing health, cognition, and lifestyle. We found that the first two profiles reflected general psychopathology (or *p factor*) associated with either reports of general poor sleep (LC1) or an absence of sleep complaints, which we defined as sleep resilience (LC2). Meanwhile, the three other profiles were driven by a specific dimension of sleep such as the use of sleep aids (LC3), sleep duration (LC4), or sleep disturbances (LC5), which were associated with distinct patterns of health, cognition, and lifestyle factors. Furthermore, identified sleep-biopsychosocial profiles displayed unique patterns of brain network organization. Our findings emphasize the crucial interplay between biopsychosocial outcomes and sleep, and the necessity to integrate sleep history to contextualize research findings and to inform clinical intake assessments ^48^.

The dominance of psychopathology markers in most of the profiles is not surprising as the RDoC framework proposed arousal and regulatory systems (i.e., circadian rhythms and sleep/wakefulness) as one of the five key domains of human functioning likely to affect mental health^49^, which is consistent with a large literature reporting significant disruption of sleep across multiple psychiatric disorders^8,50^. Although individuals with a neuropsychiatric diagnosis (e.g., schizophrenia or major depression disorder) were not included in the HCP dataset^36^, the presence of the *p* factor, defined as an individual’s susceptibility to develop any common form of psychopathology, exists on a continuum of severity and chronicity within the general population^51^. Symptoms of psychopathology mirrored each other across LC1 and LC2 but the paradoxical contrast in sleep loadings suggests that some individuals might have more resilient sleep (LC2), whereby they might be able to maintain healthy sleep patterns in the face of psychopathology. However, the cause of such resilience is unclear. Up to 80% of individuals experiencing an acute phase of mental disorder (e.g., depressive and/or anxiety episode) report sleep issues^8,52,53^, leaving a minority of individuals who do not report abnormal sleep during such episodes. The identification of LC2 supports this and suggests there might be biological or environmental protective factors in some individuals who would otherwise be considered at risk for sleep issues. However, our understanding of such protective factors is limited^54–56^. These findings also highlight the need to appreciate the complexity of psychopathology, in line with the current view that psychiatric disorders are typically comorbid and heterogeneously expressed. Nonetheless, whether this profile of sleep resilience is a stable latent component or a cross-sectional observation of fluctuating symptoms that may develop into psychopathology-related sleep complaints, needs to be further tested. Interestingly, distinctions between LC1 and LC2 were also present in the neural signatures of RSFC, which may assist in the neurobiological interpretation of the profiles. Visually inspecting LC1 and LC2 suggested an underlying increase in subcortical-cortical connectivity when sleep disturbances are associated with psychopathology. This is in alignment with the known neurophysiology of the ascending arousal system and possibly implies the existence of some level of hyperarousal in these pathways that may contribute to disturbances in sleep^57^. However, this speculation requires further targeted research to be confirmed.

Within the profiles driven by a specific sleep sub-component, LC5 also reflected some dimensions of psychopathology (i.e., anxiety, critical items and thought problems) that were only associated with the presence of global sleep disturbances. The sleep disturbance sub-component of the PSQI is broad and encompasses complaints of sleep-related breathing problems as well as multiple awakenings that could be due to nycturia, pain, nightmares, or difficulties maintaining optimal body temperature^37^. Altogether, the sleep disturbances dimension is thought to represent sleep fragmentation^58^, and thus, sleep quality. This is in line with a recent study in a large community-based cohort (i.e., UK Biobank) that found that lifetime diagnoses of psychopathology and psychiatric polygenic risk scores were more strongly associated with accelerometer-derived measures of sleep quality (i.e., fragmentation) than sleep duration per se^59^. Similarly, we found that sleep duration (driving LC4) was not associated with measures of psychopathology but rather with cognitive performance. Whether studied in the form of experimental acute sleep deprivation and chronic sleep restriction or clinical populations (e.g., insomnia with objective short sleep duration), the consequences of lack of sleep on daytime functioning and health are well-known and substantial^11,12,16,60–62^. Sleep duration affects, in varying effect sizes, both accuracy and reaction time in most cognitive tasks^11,12,61^. In our study, reports of regular short sleep duration, defined as <6-7h of total sleep time, was associated with reduced accuracy in working memory, emotional processing, language processing, delay discounting, fluid intelligence as well as longer reaction times during social cognition and emotional processing, mimicking results found in the sleep deprivation and sleep restriction literature^11,14,61–66^. Interestingly, the strong RSFC patterns associated with LC4 showed a global increase in connectivity, with localized segregation of part of the somatomotor network, which had been previously reported in neuroimaging studies of experimental acute total sleep deprivation^64,67^. Hence, this suggests that LC4 may be displaying an underlying level of sleep debt in the uncontrolled general population.

Finally, beyond sleep measures and sleep-related daytime functioning, the PSQI also evaluates the use of medication to help sleeping^37^, whether prescribed or over-the-counter (e.g., gamma-aminobutyric acid GABA_A_ receptor modulators, selective melatonin receptor agonists, selective histamine receptor antagonists, cannabinoid products, valerian)^68^. We found that LC3 was driven by the use of sleep aids and was mostly associated with reports of satisfaction in social relationships. Interestingly, while we would have expected more links between the use of sleep aids and cognitive impairment, especially in older adults^69,70^, we only found an association with visual working memory deficits but not with attentional problems. This profile specifically highlights a sub-group of young adults (22-36 years old) who experience sleep complaints and seek pharmacological solutions to manage them. As such, the associated biopsychosocial factors, in particular high sociability, could result from the effect of the drug itself on social behavior and positive mood (e.g., via potentiation of GABA transmission)^71,72^ or as a consequence of the drugs on sleep complaints^73^, which may support better emotional regulation and well-being, and consequently translate to greater satisfaction in social relationships and support systems^73,74^. We did not have information on the type nor duration of sleep aid usage as the PSQI only assesses sleep habits in the past month, which may not be a substantial period of time to observe robust changes in cognitive functioning as previously documented^70,75^ or the development of substance abuse. Indeed, while the chronic misuse of hypnotics can lead to dependence and addiction^76^, in our sample, the use of sleep aids was not associated with substance abuse.

Interestingly, alterations to the segregation/integration ratio of the somatomotor and visual cortex were common in most profiles. Highly interconnected to the whole brain, the somatomotor network is crucial for processing external stimuli and producing motor responses but is also functionally involved in bodily self-consciousness and interoception. Altered dysconnectivity patterns of the somatomotor network have been linked to variation in several domains, including general psychopathology^77,78^, cognitive dysfunction related to sleep deprivation^64^, as well as the total PSQI score^13,79^. Overall, these findings suggest that alterations to RSFC in the somatomotor network are also involved in the relationships between sleep and biopsychosocial factors and highlight the importance of understanding the role of this brain network in overall mental health and functioning.

These profiles contribute to a deeper understanding of the current debate that opposes sleep quality and sleep duration^7,80^. In line with previous studies^11,12,81^, we found that cognitive functioning was more related to sleep duration than subjective sleep quality; in addition, we found that sleep disturbances, alone (LC5) or in combination with other sleep dimensions (LC1), can be associated with the presence of psychopathology. Moreover, it is also important to note that complaints of poor sleep quality and/or short sleep duration have been both associated with increased risks of physical health outcomes and all-cause mortality^6,7^. While LC1 and LC2 presented sleep dimensions as being inextricably linked, LC3, LC4 and LC5 respectively revealed distinct facets of sleep, suggesting that while sleep dimensions are related, they can also be separable domains with specific connections to biopsychosocial factors. This is likely reflected in the finding that only LC1 and LC2 were replicable in cross-validation analyses, which may be due to LC3, LC4 and LC5 being driven by a single sleep dimension and thus, contributing only marginally to the variance. Moreover, LC1 captured a large portion of the covariance, which could be due to the presence of collinearity among the sleep and/or biopsychosocial measures. We did not apply dimensionality reduction (or any other regression techniques) to the biopsychosocial measures prior to the CCA, as the number of subjects was higher than the number of measures, and because CCA removes within-block correlations, thus allowing the amplification of specific variables that mostly characterize the correlation between sleep and biopsychosocial measures^82^. In addition, we focused on structure coefficients (not canonical coefficients) to better identify the contribution of the variables to each profile, and then computed bootstrap analyses to further assess the stability of these variables across resampling.

While unidimensional association studies have been informative, these findings reinforce the notion that sleep health is multidimensional and distinct measures of sleep quantity or quality should be considered together when investigating their influence on biopsychosocial aspects of health, cognition, and lifestyle. Future work should extend these findings and further explore the multidimensional nature of sleep health, for instance, taking into consideration the U-shaped relationship of sleep duration with biopsychosocial measures. Given the design of the PSQI, only short sleep duration (<5-6h) was considered as a sleep difficulty, neglecting the potential consequences of long sleep duration (>9h). Long sleep duration is commonly observed in hypersomnia disorders and psychopathology (e.g., schizophrenia, depression)^6,83^, as well as being associated with increased risk of cardiovascular heart disease and mortality^7,84,85^ and cognitive decline^6,20^. This U-shape observation, whereby both short and long sleep durations are associated with negative health and cognitive consequences as well as increasing markers of cerebrovascular burden (e.g., white matter hyper-intensities)^55^, may provide a window to identify mechanisms that underlie the interplay between sleep and biopsychosocial factors. Other considerations moving forward include sleep regularity and sleep timing, which are not part of the computation of the sub-components of PSQI^37^, hence their association with biopsychosocial outcomes were not investigated in this study. Furthermore, the PSQI is often interpreted with regard to its total score (combining all sub-components), which provides a binary vision of sleep quality (either good or bad sleep)^37^. In this study, we did not want to be limited by the PSQI global score but rather aimed to untangle the different dimensions (subcomponents) of sleep and their relations to biopsychosocial and neurobiological measures.

A final important distinction to be addressed is that sleep and biopsychosocial outcomes were mostly self-reported through questionnaires. Both objectively recorded and subjectively perceived estimations provide different yet meaningful information that tends to positively correlate^86^. However, it has been shown that when compared to objective estimates (i.e., polysomnography and/or actigraphy recordings), individuals with sleep complaints (i.e., chronic insomnia, obstructive sleep apnea) tend to subjectively misperceive their sleep (i.e., duration, sleep latency)^25,26,87,88^. The degree of discrepancy between objective and subjective measures (i.e., sleep state misperception) has been correlated with worse sleep quality^89,90^ as well as compromised reports of daytime functioning^24^. While objective measurements might have exposed divergent associations between sleep and biopsychosocial factors, the profiles reported here arguably support greater clinical validity, where the subjective complaints are often what drives an individual to seek out healthcare. Our study emphasizes that considering individuals’ sleep experience can support clinicians to make more accurate initial assessments and navigate the course of treatment and interventions.

The awareness and interest surrounding sleep as a crucial pillar of health is growing rapidly^91^. However, the role of sleep in general health is complex, multifaceted, and largely unknown. The multidimensional approach applied in this large sample of healthy young adults is a first step that we argue should be implemented in future research incorporating sleep. We highlight the observation of five distinct sleep patterns associated with specific combinations of biological, psychological and socio-environmental factors. These findings support that sleep is emerging as a distinguishable factor that can assist in disentangling the complex heterogeneity of human health. As the capacity for large-scale human research continues to grow, integrating sleep dimensions at such a scale is not only feasible in terms of evaluation, but presents a unique opportunity for translational application. Sleep is a modifiable lifestyle factor and can be investigated in model organisms as well as in humans, and as such is well positioned to identify potential converging mechanisms and intervention pathways or tools. The current study emphasizes that by using a multidimensional approach to identify distinct sleep-biopsychosocial profiles we can begin to untangle the interplay between individuals’ variability in sleep, health, cognition, lifestyle, and behavior—equipping research and clinical settings to better support individuals’ well-being.

## METHODS

### Participants

Data for this study were obtained from the S1200 release of the publicly available Human Connectome Project (HCP) dataset^36^. The HCP dataset comprises multimodal MRI data, including structural MRI, diffusion MRI, resting-state and task functional MRI (fMRI) data, as well as a broad range of behavioral measures collected in young healthy subjects (aged 22-36). Details about imaging acquisition parameters and data collection^36^ as well as the list of available behavioral and demographics measures (HCP S1200 Data Dictionary)^92^ can be found elsewhere. Of note, the HCP dataset comprises a large number of related individuals (i.e., siblings and twins). Of the 1,206 total subjects available from the HCP S1200 release, we excluded 403 participants with missing/incomplete data, and 33 participants with visual impairment that might have impacted their task performance in the scanner. Our final sample comprised 770 participants (53.76% female, 28.86 ± 3.61 years old). We decided to keep participants (N=94) who tested positive for any substance (including alcohol, marijuana, and other drugs) on the day of the MRI, as substance use has intricate links to sleep, and we did not want to exclude the possibility of finding potential substance use-related sleep profiles. However, we also re-computed our analyses after excluding these individuals (N=676) and found very similar results (see Table S5). Out of these 770 participants, 723 passed MRI quality control and were included in the *posthoc* RSFC analyses.

### Sleep assessment

Participants were administered the Pittsburgh Sleep Quality Index^37^ (PSQI) to assess different aspects of their sleep over the past month. To define sleep in our study, we used the 7 sub-components of the PSQI which characterize different sleep dimensions, namely (i) sleep satisfaction, (ii) sleep latency, (iii) sleep duration, (iv) sleep efficiency, (v) sleep disturbance, (vi) sleep medication, and (vii) daytime functioning. Sub-components are calculated through 4 questions on the timing of sleep habits and 6 Likert-scale questions from 0 to 3, 0 being best and 3 being worst.

### Biopsychosocial assessment

118 biopsychosocial measures were selected from the HCP dataset (see complete list in Table S1). These measures included self-reported assessments of current and past mental health and substance use, questionnaires on personality, affect, lifestyle and demographics, cognitive tasks tapping on different processes such as working memory or social cognition performed either inside or outside the MRI, and physical assessments (e.g., blood pressure). These measures did not undergo any dimensionality reduction or clustering by biopsychosocial domain in order to preserve granularity in the way they would be associated with sleep dimensions. Biopsychosocial measures with large amounts of missing data were excluded, as well as similar measures that were likely to be redundant. Biopsychosocial measures were categorized by behavioral domain (e.g., cognition, physical health) based on the way they had been described in the HCP dataset^36,92^.

### Canonical correlation analysis

Canonical Correlation Analysis (CCA)^93,94^, a multivariate data-driven approach, was applied to the sleep and biopsychosocial measures. CCA derives latent components (LCs, i.e., canonical variates), which are optimal linear combinations of the original data, by maximizing *correlation* between two data matrices (i.e., sleep and biopsychosocial measures). The rank of the correlation matrix determines the number of derived LCs (i.e., in this case the number of sleep measures, hence 7 LCs). Each sleep-biopsychosocial LC is characterized by a pattern of sleep weights and a corresponding pattern of biopsychosocial weights (i.e., canonical coefficients). Linear projection of sleep (or biopsychosocial) data onto sleep (or biopsychosocial) weights yielded participant-specific composite scores for sleep (or biopsychosocial) measures (i.e., canonical scores). The contribution of original sleep and biopsychosocial loadings to each LC was determined by computing Pearson’s correlations between sleep (or biopsychosocial) data and participant-specific scores for sleep (or biopsychosocial factors) to obtain sleep and biopsychosocial *loadings* (i.e., canonical structure coefficients)^95,96^. Canonical structure coefficients reflect the direct contribution of a predictor (e.g., one sleep dimension) to the predictor criterion (e.g., LC1) independently of other predictors (e.g., LCs 2-7), which can be critical when predictors are highly correlated between each other (i.e., in presence of multicollinearity)^97^. We did not employ dimensionality reduction (e.g., via principal components analysis), as the sample size (N=770) exceeded the number of sleep (7 measures) and biopsychosocial measures (118 measures) being modelled. Statistical significance of each of the 7 LCs was determined by permutation testing (10,000 permutations) followed by FDR correction. Given the high prevalence of related participants in the HCP dataset, family structure was maintained during permutations (using the PALM package^98,99^), whereby monozygotic twins, dizygotic twins, and non-twin siblings were only permuted within their respective groups. Finally, the loadings’ stability was determined using bootstrap resampling to estimate confidence intervals for the loadings, by deriving 1,000 samples with replacement from participants’ sleep and biopsychosocial data.

### MRI acquisition and processing

All imaging data were acquired on a customized Siemens 3T Skyra scanner at Washington University (St Louis, MI). Four runs of resting-state fMRI were collected over two sessions across two separate days. Each run included 1,200 frames using a multi-band sequence at 2-mm isotropic spatial resolution with a TR of 0.72 s for 14.4 minutes. The structural images were acquired at 0.7-mm isotropic resolution. Further details of the data collection and HCP preprocessing are available elsewhere^36,100,101^. Notably, cortical and subcortical data underwent ICA-FIX^102,103^ and were saved in the CIFTI gray ordinate format. The surface (fs_LR) data were aligned with MSM-All^104^. As ICA-FIX does not fully eliminate global motion-related and respiratory-related artifacts^105,106^, additional censoring and nuisance regression were performed^107,108^. In particular, volumes with framewise displacement (FD) > 0.2mm, and root-mean-square of voxel-wise differentiated signal (DVARS) > 75 were marked as outliers and censored, along with one frame before and two frames after the outlier volume^109,110^. Any uncensored segment of data that lasted fewer than five contiguous volumes was also excluded from analysis, as well as runs with >50% censored frames. Additionally, the global signal obtained by averaging signal across all cortical vertices and its temporal derivatives (ignoring censored frames) were also regressed out from the data because previous studies have suggested that global signal regression strengthens the association between RSFC and behavioral traits^107^. As there is ongoing debate on the use of global signal regression (GSR) as a means of fMRI preprocessing^107,111–113^, additional reliability analysis was performed on data preprocessed using a component-based noise correction method (CompCor)^47^ instead of GSR.

RSFC was computed among 400 cortical parcels^46^ and 19 subcortical regions^45^ using Pearson’s correlation (excluding the censored volumes). The subcortical regions were in subject-specific volumetric space as defined by FreeSurfer^45^, and comprised the left and right cerebellum, thalamus, caudate, putamen, pallidum, hippocampus, accumbens, amygdala, ventral diencephalon, and brainstem. For each participant, RSFC was computed for each run, Fisher z-transformed, and then averaged across runs and sessions, yielding a final 419 x 419 RSFC matrix for each participant.

### RSFC analyses

To investigate the neurobiological substrates of the sleep-biopsychosocial profiles derived in the CCA, we computed generalized linear models (GLM) between participant’s canonical scores (i.e., averaged sleep and biopsychosocial scores) and their RSFC data. Age, sex, and level of education were first regressed out from the RSFC data.

To obtain an analysis at the large-scale network level and limit the number of multiple comparisons, we computed a network-wise GLM, whereby the whole-brain RSFC data was averaged within and between the 17 large-scale brain networks^46^ and subcortical regions^45^, resulting in 18 x 18 RSFC matrices. Next, we applied a GLM for each network edge (i.e., average connectivity between two brain networks), with participants’ component-specific canonical scores as the predictor and RSFC edge as the response. Each GLM yielded a beta coefficient and associated *T* statistic, as well as an *F* statistic and associated *p* value obtained from a hypothesis test that all coefficient estimates were equal to zero. Statistical significance for each RSFC network edge was determined by applying FDR correction (*q* < 0.05) on all *p* values (along with other posthoc analyses). For a more granular view, we also computed a GLM for each RSFC edge (i.e., connectivity between two brain regions) using whole-brain RSFC between all 419 brain regions. For a complete view of the component-specific RSFC signatures, we plotted both the uncorrected region-wise GLM beta coefficients (e.g., Figure 2C) and FDR-corrected network-wise GLM beta coefficients (e.g., Figure 2D).

Measures of integration and segregation were computed on the GLM beta coefficient connectivity matrix associated with each LC using functions from the Brain Connectivity Toolbox^114^. Firstly, the input-weighted connection matrix was normalized. Next, each 419 cortical parcel was assigned to one of the 7 large-scale brain networks and subcortical regions ^44^. Within-network connectivity was estimated by calculating the module-degree Z score (within-module strength) for each region. The extent to which a parcel connects across all networks was quantified using the participation coefficient, (between-module strength). For each cortical parcel, the ratio of normalized within:between module strength values was calculated and interpreted as a measure for the balance of integration and segregation of functional brain connectivity^115^. Nodes with high within-but low between-module strength are likely to facilitate network segregation, while nodes with higher between-module strength (i.e., connector hubs) are likely to facilitate global integration^114^.

### Control analyses

We ran several control analyses to evaluate the robustness of our findings. First, we applied 5-fold cross-validation (accounting for family structure) to assess the generalizability of our sleep-biopsychosocial profiles by training a CCA model on 80% of the data and testing it on the remaining 20% of the data. For each fold, we projected the sleep and biopsychosocial canonical coefficients of the training data on the sleep and biopsychosocial data of the test data, to obtain sleep and biopsychosocial scores, and computed Pearson’s correlations between these scores. Second, we evaluated the impact of the covariates on our profiles as well as the impact of other potential confounds, including race, ethnicity, and familial psychiatric history. Third, we re-computed the CCA analysis after excluding participants who had tested positive for any substance use on the day of the MRI. Fourth, we re-computed the CCA analysis after excluding physical health (i.e., body mass index, hematocrit, blood pressure) and sociodemographic (i.e., employment status, household income, in-school, relationship status) variables from the biopsychosocial matrix. Fifth, to mitigate scale magnitude discrepancies between different measures, we re-computed the CCA analysis after applying quantile normalization on sleep and biopsychosocial measures. We also assessed the robustness of our imaging results in several ways. As GSR is a controversial preprocessing step^107,112,113^, we re-computed the GLM analysis using RSFC data that underwent CompCor^47^ instead of GSR. Some subjects were noticed to have likely fallen asleep during scanning (list not publicly available^116^). As a first step, we re-computed the GLM after excluding these subjects (N=100); next, we sought to determine whether these participants scored high on any of the profiles, by comparing their sleep/biopsychosocial composite scores with awake participants using t-tests. We re-computed the GLM analyses by using sleep and biopsychosocial canonical scores instead of averaged scores. Finally, integration and segregation measures were also computed on the average RSFC matrix of the whole sample. FDR correction (*q* < 0.05) was applied to all posthoc tests.

## Data and code availability

Data from the HCP dataset is publicly available (https://www.humanconnectome.org/). The brain parcellation can be obtained here (https://github.com/ThomasYeoLab/CBIG/tree/master/stable_projects/brain_parcellation/Schaefer20 18_LocalGlobal), while the code for the CCA analysis and figures can be found here (https://github.com/valkebets/sleep_biopsychosocial_profiles). Chord diagrams were generated using previously published code (https://github.com/ThomasYeoLab/CBIG/tree/master/stable_projects/predict_phenotypes/ChenTam 2022_TRBPC/figure_utilities/chord).

## ACKNOWLEDGEMENT

Any opinions, findings, conclusions or recommendations expressed in this material are those of the authors and do not reflect the views of the Singapore NRF, Singapore NMRC, MOH or Temasek Foundation. Our research also utilized resources provided by the Center for Functional Neuroimaging Technologies, P41EB015896 and instruments supported by 1S10RR023401, 1S10RR019307, and 1S10RR023043 from the Athinoula A. Martinos Center for Biomedical Imaging at the Massachusetts General Hospital. The computational work was partially performed using resources of the National Supercomputing Centre, Singapore (http://www.nscc.sg). Data were provided by the Human Connectome Project, WU-Minn Consortium (Principal Investigators: David Van Essen and Kamil Ugurbil; 1U54MH091657) funded by the 16 NIH Institutes and Centers that support the NIH Blueprint for Neuroscience Research; and by the McDonnell Center for Systems Neuroscience at Washington University. AAP has been supported by the American Academy of Sleep Medicine (AASM), Fondation Lemaire and fellowships from Concordia University, Centre de Recherche de l’Institut Universitaire de Gériatrie de Montréal (CRIUGM) and PERFORM Center. VK has been supported by the Transforming Autism Care Consortium and the Montreal Neurological Institute. NC has been supported by the Fonds de Recherche du Québec – Santé and a fellowship from the CRIUGM. TDV is currently supported by CIHR grants, the Natural Sciences and Engineering Research Council of Canada, the Canada Foundation for Innovation and the Fonds de Recherche du Québec – Santé. BTTY is currently supported by the NUS Yong Loo Lin School of Medicine (NUHSRO/2020/124/TMR/LOA), the Singapore National Medical Research Council (NMRC) LCG (OFLCG19May-0035), NMRC CTG-IIT (CTGIIT23jan-0001), NMRC STaR (STaR20nov-0003), Singapore Ministry of Health (MOH) Centre Grant (CG21APR1009), the Temasek Foundation (TF2223-IMH-01), and the United States National Institutes of Health (R01MH120080 & R01MH133334). Finally, we thank Dr. Joshua Gooley for his helpful comments in the previous versions of the work.

## AUTHOR CONTRIBUTION STATEMENT

Conceptualization: NMYK, BTTY, VK, AAP; Data curation: VK, NMYK, JL, AAP; Formal analysis: VK, NMYK, NEC, AAP; Methodology: VK, NMYK, NEC; Visualization: AAP, VK, NEC; Interpretation: AAP, VK, NEC, RT; Writing - original draft: AAP, VK, NMYK; editing and reviewing: AAP, VK, RT, NEC, BTTY, FBP, TTDV, JL, MWLC, NMYK.

