## Supplemental Material for "A multidimensional investigation of sleep and biopsychosocial profiles with associated neural signatures"

Title:

##### Canonical correlation analysis

Canonical Correlation Analysis (CCA) is a multivariate data-driven approach that derives latent components (LCs; i.e., canonical variates), that maximize *correlation* between two set of multidimensional variables (*1*, *2*). We applied the *canoncorr* function from Matlab 2018b to our dataset, to obtain latent components that are optimal linear combinations of the sleep-biopsychosocial data.

The CCA analysis was computed as follows. Sleep and biopsychosocial measures are stored in matrices $X$ (770 × 7) and $Y$ (770 × 118). First, $X$ and $Y$ each undergo orthogonal decomposition such that:

$$X = Q1 \times R1$$
$$Y = Q2 \times R2$$

where $Q1$ and $Q2$ are orthogonal matrices, and $R1$ and $R2$ are upper unitary matrices. Orthogonal matrices are then multiplied to obtain a correlation matrix:

$$Q = Q1^T \times Q2$$

Onto which singular value decomposition (SVD) is applied:

$$Q = A \times S \times B^T$$

This results in two singular vector matrices, $A$ and $B$, and a diagonal matrix containing the singular values, $S$. The singular vector matrices of each LC form the sleep weights (7 × 7), and biopsychosocial weights (118 × 7). When $A$ and $B$ are linearly projected onto respective sleep and biopsychosocial scores, $X$ and $Y$, it yields maximally correlated canonical variates:

$$U \ = X \times A$$
$$V \ = Y \times B$$

These canonical scores (composite scores) can be interpreted as participants' loadings on the sleep and biopsychosocial weights of each component. The contribution of original sleep and biopsychosocial loadings to each LC was determined by computing Pearson's correlations between sleep (or biopsychosocial) data and participant-specific scores for sleep (or biopsychosocial) to obtain sleep and biopsychosocial *loadings* (i.e., canonical structure coefficients)(*3*, *4*). Canonical structure coefficients reflect the direct contribution of a predictor to the predictor criterion independently of other predictors, which can be critical when predictors are highly correlated with each other (i.e., in the presence of multicollinearity)(*5*). Statistical significance of each LC was determined by permutation testing (10,000 permutations) followed by FDR correction. Given the high prevalence of related participants in the HCP dataset for family structure was maintained during permutations (using the PALMS package(*6*, *7*), whereby monozygotic twins, dizygotic twins, and non-twin siblings were only permuted within their respective groups. Finally, the loadings' stability was determined using bootstrap resampling to estimate confidence intervals for the loadings, by deriving 1,000 samples with replacement from participants' sleep and biopsychosocial data.

#### Supplementary Results

##### Post-hoc associations with socio-demographics, health, and family history of mental health

We found several significant associations between LC composite scores and biopsychosocial factors (**Table S4**). In LC1, we found a significantly negative correlation between higher sleep and biopsychosocial composite scores and education level, and significant differences in sleep and biopsychosocial composite scores between household income categories (i.e., generally higher scores in those living in a low-income household). Biopsychosocial

composite scores were also higher in Hispanic/Latino participants and in participants with a paternal history of depression.
In LC2, we found a significantly positive correlation between higher sleep and biopsychosocial composite scores and education level; significantly higher biopsychosocial composite scores in participants from lower household income); lower composite sleep scores in participants working full-time vs. part-time vs. those not working; higher sleep scores in those not in school; higher sleep composite scores in participants with a paternal history of depression, higher biopsychosocial composite scores in those with a maternal history of anxiety or a paternal history of addiction.
In LC3, we found significant differences in sleep composite scores between participants from different racial backgrounds (i.e., higher scores in participants who identified as White), and a significant negative correlation between sleep composite scores and BMI.
In LC4, we found a significant association between higher sleep and biopsychosocial composite scores and higher age as well as lower education level, and between higher sleep scores and higher BMI. There were also significant differences in sleep composite scores between participants from different racial backgrounds. We also found higher biopsychosocial composite scores in participants working full time but also in those not in school. Finally, we find lower biopsychosocial scores in participants with a paternal history of anxiety.
Finally in LC5, we found higher sleep and biopsychosocial composite scores in female participants; a significant association between higher sleep and biopsychosocial composite scores and lower education level; and significant differences between participants from different racial backgrounds.

#### Control RSFC analyses

We computed some control GLM analyses with RSFC data (**Figure S3**).
First, we computed a GLM using RSFC data that underwent CompCor instead of global signal regression that showed high correlations with the main GLM analysis (LC1: r=0.75; LC2: r=0.76; LC3: r=0.78; LC4: r=0.51; LC5: r=0.77).
Next, we computed a GLM analysis after excluding subjects that likely fell asleep in the scanner (N=100), which showed high correlations with the main GLM analysis (LC1: r=0.90; LC2: r=0.87; LC3: r=0.95; LC4: r=0.95; LC5: r=0.95).
Third, we computed a GLM analysis between RSFC and *sleep* composite scores (instead of averaged composite scores), which showed moderate to high correlations with the main GLM analysis (LC1: r=0.72; LC2: r=0.60; LC3: r=0.74; LC4: r=0.52; LC5: r=0.66).
We did the same between RSFC and *biopsychosocial* composite scores (instead of averaged composite scores), and found moderate to high correlations with the main GLM analysis (LC1: r=0.69; LC2: r=0.62; LC3: r=0.63; LC4: r=0.46; LC5: r=0.67).
Finally, we computed integration and segregation measures on the average RSFC matrix of the whole sample to compare our findings per LC with the average (**Figure S5**).

#### Control analyses with asleep participants

We computed post-hoc t-tests to assess differences in sleep (or biopsychosocial) composite scores between participants who likely stayed awake (N=623) in the scanner vs. those who likely fell asleep in the scanner (N=100). Participants that likely fell asleep in the scanner had significantly higher biopsychosocial composite scores on LC1 ($t$=2.93, $p$=0.003), significantly lower sleep composite scores on LC3 ($t$=-2.42, $p$=0.016), and significantly higher sleep ($t$=2.25, $p$=0.025) and biopsychosocial composite scores on LC4 ($t$=2.85, $p$=0.005) compared to those that likely stayed awake in the scanner (**Figure S4).**
When inspecting the distribution of sleep and biopsychosocial composite scores of these participants, they did not appear to be driving any of the LCs (**Figure S6**).

### Supplementary Tables and Figures

**Table S1**. Sleep and non-sleep biopsychosocial measures used in the CCA (indicated in black), or in post-hoc analyses (indicated in blue), or in both (indicated in green).

| **Domain** | **Scale** | **Subdomain** | **Measure** |
|---|---|---|---|
| Sleep | Pittsburgh Sleep Quality Index (*8*) | Component 1: Sleep satisfaction | |
| | | Component 2: Sleep latency | |
| | | Component 3: Sleep duration | |
| | | Component 4: Sleep efficiency | |
| | | Component 5: Sleep disturbance | |
| | | Component 6: Sleep meds | |
| | | Component 7: Daytime functioning | |
| Socio-demographics | | | Age |
| | | | Sex |
| | | | Education level (years) |
| | | | Race |
| | | | Ethnicity |
| | | | Employment status |
| | | | Household income |
| | | | School status |
| | | | Relationship status |
| Physical health | | Physical function | Body mass index (BMI) |
| | | | Blood pressure* |
| | | | Hematocrit levels** |
| Cognition (outside the scanner) | NIH Toolbox (https://www.nihtoolbox.org/) | Episodic memory | Picture sequence memory test |
| | | Executive function, cognitive flexibility | Dimensional change card sort test |
| | | Executive function, inhibition | Flanker inhibitory control and attention test |
| | | Language, reading decoding | Oral reading recognition test |
| | | Language, vocabulary comprehension | Picture vocabulary test |
| | | Processing speed | Pattern comparison processing speed test |
| | | Working memory | List sorting working memory test |
| | Penn Progressive matrices (*9*) | Fluid intelligence | Accuracy |
| | | | RT |
| | Variable Short Penn Line Orientation Test (*10*, *11*) | Visual-spatial processing | Accuracy |
| | | | RT |
| | Short Penn Continuous Performance Test (Number/Letter Version) (*10–12*) | Sustained attention | True positives |
| | | | Sensitivity |
| | | | Specificity |
| | | | Accuracy |

| | | | |
|---|---|---|---|
| | Penn Word Memory Test, Form A (*10*, *11*) | Verbal episodic memory | RT |
| | Penn Emotion Recognition Test (*10*, *11*) | Emotion processing | Accuracy- Anger |
| | | | Accuracy- Fear |
| | | | Accuracy- Happy accuracy |
| | | | Accuracy- Neutral accuracy |
| | | | Accuracy- Sad accuracy |
| | | | Accuracy- Total |
| | | | RT- Total |
| | Delay discounting task (*13*, *14*) | Self-regulation, impulsivity | Area under the curve (AUC) for $200 |
| | | | Area under the curve (AUC) for $40,000 |
| Cognition (in-scanner) | Language task (*15*) | Language processing | Accuracy |
| | | | RT |
| | Relational task (*16*) | Relational processing | Accuracy |
| | | | RT |
| | Social task (*17*, *18*) | Social cognition, theory of mind | Accuracy random stimuli rated random |
| | | | RT random stimuli rated random |
| | | | Accuracy social stimuli rated social |
| | | | RT social stimuli rated social |
| | Working memory n-back task (*19–29*) | Working memory, cognitive control | Accuracy |
| | | | RT |
| | Gambling task (*30*) | Incentive processing | Accuracy smaller prediction |
| | | | RT smaller prediction |
| | | | Accuracy larger prediction |
| | | | RT larger prediction |
| | Hariri Emotion task (*31*) | Emotional processing | Accuracy |
| | | | RT |
| Affect | NIH Toolbox (https://www.nihtoolbox.org/) | Negative affect | Anger- affect |
| | | | Anger- hostility |
| | | | Anger- physical aggression |
| | | | Fear- affect |
| | | | Fear- somatic arousal |
| | | | Sadness |
| | | Psychological well-being | Positive affect |
| | | | General life satisfaction |
| | | | Meaning and purpose |
| | | Social relationships | Emotional support |
| | | | Instrumental support |
| | | | Friendship |
| | | | Loneliness |
| | | | Perceived hostility |
| | | | Perceived rejection |
| | | Stress and self-efficacy | Perceived stress |
| | | | Self-efficacy |
| Sensory | NIH Toolbox (https://www.nihtoolbox.org/) | Sensory processing | Pain interference |
| Personality | Costa and McRae Neuroticism/Extroversion/Openness Five Factor Inventory (NEO-FFI) (*32*) | Personality traits | Agreeableness |
| | | | Openness to experience |
| | | | Conscientiousness |
| | | | Neuroticism |

| | | | |
|---|---|---|---|
| | | | Extroversion/introversion |
| Mental health | Achenbach Adult Self-Report (ASR) (*33*) | Life function, psychiatric clinical symptoms | Anxious depressed |
| | | | Withdrawn |
| | | | Somatic complaints |
| | | | Thought problems |
| | | | Rule breaking behavior |
| | | | Aggressive behavior |
| | | | Attention problems |
| | | | Intrusive problems |
| | | | Other problems |
| | | | Internalizing problems |
| | | | Externalizing problems |
| | | | Total problems |
| | | | Critical items |
| | | | DSM Anxiety problems |
| | | | DSM ADHD problems |
| | | | DSM Hyperactivity problems |
| | | | DSM Depressive problems |
| | | | DSM Avoidant personality problems |
| | | | DSM Antisocial personality problems |
| | | | DSM Somatic problems |
| | | | DSM Inattention problems |
| | Semi-Structured Assessment for the Genetics of Alcoholism (SSAGA) (*34*) | Psychiatric clinical symptoms | Agoraphobia |
| | | | Childhood conduct |
| | | | Major depressive episodes |
| | | | Number of depressive symptoms |
| | | | Panic disorder |
| | | Family history of psychiatric and neurological disorders | Maternal history of schizophrenia or psychosis |
| | | | Paternal history of schizophrenia or psychosis |
| | | | Maternal history of depression |
| | | | Paternal history of depression |
| | | | Maternal history of bipolar disorder |
| | | | Paternal history of bipolar disorder |
| | | | Maternal history of anxiety |
| | | | Paternal history of anxiety |
| | | | Maternal history of drug or alcohol problems |
| | | | Paternal history of drug or alcohol problems |
| | | | Maternal history of Alzheimer's disease or dementia |
| | | | Paternal history of Alzheimer's disease or dementia |
| | | | Maternal history of Parkinson's disease |
| | | | Paternal history of Parkinson's disease |
| | | | Maternal history of Tourette's syndrome |
| | | | Paternal history of Tourette's syndrome |
| Substance use | Semi-Structured Assessment for the Genetics of Alcoholism (SSAGA) (*34*) | Substance use, abuse and dependence | Alcohol age first use |
| | | | Drinks per day past 12 months |
| | | | Frequency alcohol use past 12 months |
| | | | Frequency drunk past 12 months |
| | | | Drinks per day heaviest period |
| | | | Frequency alcohol use heaviest period |

| | | | |
|---|---|---|---|
| | | | Frequency drunk heaviest period |
| | | | Lifetime alcohol abuse diagnosis |
| | | | Lifetime alcohol abuse symptoms |
| | | | Lifetime alcohol dependence diagnosis |
| | | | Lifetime alcohol dependence symptoms |
| | | | Times used cocaine |
| | | | Times used hallucinogens |
| | | | Times used marijuana |
| | | | Times used nonmarijuana illicit drugs |
| | | | Times used opiates |
| | | | Times used sedatives |
| | | | Times used stimulants |
| | | | Marijuana dependence |
| | Fagerstrom Test for Nicotine Dependence (*35*, *36*) | Nicotine dependence | History of smoking |
| | | | Currently smoking |

**Blood pressure was categorized into (i) hypotension (systolic blood pressure (BP) ≤ 90 and/or diastolic BP ≤ 60), (ii) normal BP (systolic BP < 120 and diastolic BP = 80), (iii) elevated BP (systolic BP between 120-129 and/or diastolic BP < 80), (iv) hypertension stage 1 (systolic BP between 130-139 and/or diastolic BP between 80-89), and (v) hypertension stage 2 (systolic BP > 140 and/or diastolic BP ≥ 90).*

*** Hematocrit levels were averaged across the day 1 and day 2 measurements.*

**Table S2**. CCA loadings and z-scores for sleep and biopsychosocial measures for LCs 1-5. Loadings with significant bootstrapped z-scores that survived FDR correction ($q < 0.05$) are indicated in bold.

| Variable | LC1 loading (z-score) | LC2 loading (z-score) | LC3 loading (z-score) | LC4 loading (z-score) | LC5 loading (z-score) |
|---|---|---|---|---|---|
| Comp1 Sleep satisfaction (PSQI) | **-0.72 (-4.06)** | 0.22 (1.19) | -0.23 (-1.14) | 0.21 (0.91) | **0.44 (2.37)** |
| Comp2 Sleep latency (PSQI) | **-0.72 (-3.88)** | **0.35 (2.09)** | 0.21 (0.97) | -0.16 (-0.75) | -0.06 (-0.30) |
| Comp3 Sleep duration (PSQI) | -0.22 (-1.00) | 0.13 (0.51) | -0.20 (-0.72) | **0.83 (3.86)** | -0.14 (-0.43) |
| Comp4 Sleep efficiency (PSQI) | -0.33 (-1.32) | 0.22 (1.25) | -0.18 (-0.83) | -0.15 (-0.54) | -0.18 (-0.58) |
| Comp5 Sleep disturbance (PSQI) | **-0.57 (-3.23)** | 0.17 (0.90) | -0.16 (-0.75) | -0.07 (-0.26) | **-0.64 (-3.57)** |
| Comp6 Sleep meds (PSQI) | **-0.39 (-2.09)** | -0.17 (-0.55) | **0.85 (5.71)** | 0.13 (0.44) | -0.06 (-0.26) |
| Comp7 Daytime functioning (PSQI) | **-0.57 (-3.21)** | **-0.74 (-4.57)** | -0.33 (-1.19) | 0.02 (0.08) | -0.03 (-0.18) |
| Employment status | **0.10 (1.81)** | -0.02 (-0.30) | -0.03 (-0.37) | **0.14 (1.87)** | -0.01 (-0.10) |
| Household income | **0.13 (1.73)** | **0.14 (1.67)** | -0.04 (-0.42) | 0.14 (1.41) | **0.23 (2.73)** |
| School status | 0.00 (-0.05) | **-0.17 (-2.35)** | 0.06 (0.59) | **0.16 (1.89)** | -0.01 (-0.14) |
| Relationship status | 0.02 (0.42) | 0.00 (0.03) | -0.11 (-1.41) | 0.04 (0.47) | 0.11 (1.25) |
| BMI | -0.07 (-0.91) | 0.10 (1.04) | **-0.23 (-2.72)** | 0.11 (1.07) | -0.15 (-1.56) |
| Hematocrit | **-0.09 (-1.76)** | 0.06 (1.09) | -0.04 (-0.53) | -0.06 (-0.88) | 0.07 (1.11) |
| Blood pressure | 0.02 (0.31) | 0.11 (1.27) | -0.09 (-0.89) | -0.13 (-1.20) | **-0.23 (-2.19)** |
| Anxious depressed (ASR) | **-0.47 (-3.69)** | **-0.32 (-3.79)** | -0.07 (-0.56) | 0.04 (0.38) | -0.15 (-1.53) |
| Withdrawn (ASR) | **-0.30 (-3.00)** | **-0.21 (-2.30)** | -0.15 (-1.47) | 0.02 (0.22) | -0.12 (-1.17) |
| Somatic complaints (ASR) | **-0.62 (-3.83)** | -0.11 (-1.24) | 0.03 (0.32) | -0.01 (-0.11) | **-0.16 (-2.12)** |
| Thought problems (ASR) | **-0.38 (-3.27)** | **-0.31 (-3.05)** | -0.17 (-1.36) | 0.11 (0.94) | **-0.20 (-2.01)** |
| Rule breaking behavior (ASR) | **-0.30 (-3.16)** | **-0.17 (-2.11)** | -0.12 (-1.29) | 0.04 (0.40) | -0.09 (-1.13) |
| Aggressive behavior (ASR) | **-0.35 (-2.91)** | **-0.21 (-2.47)** | -0.05 (-0.47) | 0.16 (1.54) | **-0.20 (-2.23)** |

| | | | | | |
|---|---|---|---|---|---|
| Attention problems (ASR) | **-0.41 (-3.65)** | **-0.45 (-4.17)** | -0.24 (-1.60) | -0.05 (-0.36) | -0.02 (-0.22) |
| Intrusive problems (ASR) | -0.10 (-1.62) | **-0.12 (-1.75)** | -0.10 (-1.33) | 0.01 (0.17) | -0.05 (-0.63) |
| Other problems (ASR) | **-0.48 (-3.57)** | **-0.34 (-3.82)** | -0.09 (-0.68) | 0.05 (0.40) | **-0.18 (-1.86)** |
| Internalizing problems (ASR) | **-0.56 (-3.76)** | **-0.28 (-3.14)** | -0.07 (-0.58) | 0.03 (0.25) | **-0.17 (-1.83)** |
| Externalizing problems (ASR) | **-0.34 (-3.04)** | **-0.22 (-2.60)** | -0.11 (-1.04) | 0.10 (1.02) | **-0.16 (-1.83)** |
| Total problems (ASR) | **-0.54 (-3.63)** | **-0.36 (-3.70)** | -0.14 (-1.03) | 0.05 (0.43) | **-0.17 (-1.76)** |
| Critical items (ASR) | **-0.44 (-3.27)** | **-0.40 (-4.37)** | -0.06 (-0.41) | 0.08 (0.64) | **-0.22 (-2.17)** |
| DSM Anxiety problems (ASR) | **-0.50 (-3.44)** | **-0.27 (-3.02)** | -0.08 (-0.65) | 0.12 (1.10) | **-0.18 (-1.93)** |
| DSM ADHD problems (ASR) | **-0.40 (-3.51)** | **-0.37 (-4.54)** | -0.08 (-0.60) | 0.03 (0.30) | -0.06 (-0.65) |
| DSM Hyperactivity problems (ASR) | **-0.39 (-3.34)** | **-0.21 (-2.63)** | 0.09 (0.91) | 0.05 (0.51) | -0.12 (-1.48) |
| DSM Depressive problems (ASR) | **-0.61 (-3.99)** | **-0.32 (-3.47)** | -0.10 (-0.82) | -0.05 (-0.45) | -0.05 (-0.58) |
| DSM Avoidant personality problems (ASR) | **-0.26 (-3.18)** | **-0.20 (-2.31)** | -0.15 (-1.45) | -0.03 (-0.24) | -0.07 (-0.68) |
| DSM Antisocial personality problems (ASR) | **-0.30 (-2.79)** | **-0.18 (-2.13)** | -0.14 (-1.51) | 0.06 (0.59) | -0.12 (-1.35) |
| DSM Somatic problems (ASR) | **-0.42 (-3.45)** | **-0.17 (-2.08)** | 0.06 (0.64) | -0.01 (-0.07) | **-0.19 (-2.35)** |
| DSM Inattention problems (ASR) | **-0.31 (-3.32)** | **-0.42 (-4.40)** | -0.20 (-1.42) | 0.01 (0.09) | 0.00 (0.03) |
| Childhood conduct (SSAGA) | **-0.23 (-3.07)** | -0.05 (-0.55) | **-0.20 (-2.93)** | 0.00 (0.03) | -0.10 (-1.21) |
| Panic disorder (SSAGA) | **-0.14 (-2.23)** | -0.04 (-0.58) | -0.01 (-0.12) | -0.04 (-0.57) | 0.02 (0.23) |
| Agoraphobia (SSAGA) | -0.08 (-1.57) | -0.11 (-1.28) | **-0.20 (-2.78)** | -0.01 (-0.08) | 0.02 (0.26) |
| Major depressive episodes (SSAGA) | **-0.33 (-3.51)** | **-0.23 (-3.09)** | 0.03 (0.25) | -0.03 (-0.29) | -0.07 (-0.81) |
| Number of depressive symptoms (SSAGA) | **-0.35 (-3.49)** | **-0.23 (-3.22)** | 0.01 (0.12) | -0.02 (-0.23) | -0.04 (-0.45) |
| Picture sequence memory accuracy (NIH) | 0.00 (0.07) | 0.04 (0.43) | **-0.22 (-2.81)** | **-0.16 (-1.74)** | 0.02 (0.24) |
| Dimensional change card sort accuracy (NIH) | 0.00 (-0.07) | 0.09 (1.46) | 0.01 (0.07) | 0.01 (0.09) | 0.07 (0.88) |
| Flanker inhibitory control attention accuracy (NIH) | 0.02 (0.41) | 0.10 (1.56) | -0.06 (-0.88) | 0.07 (0.92) | 0.03 (0.44) |
| Oral reading recognition accuracy (NIH) | -0.02 (-0.36) | **-0.20 (-2.70)** | 0.03 (0.29) | -0.14 (-1.47) | **0.19 (2.13)** |
| Picture vocabulary accuracy (NIH) | -0.04 (-0.69) | **-0.13 (-1.97)** | 0.00 (0.00) | 0.03 (0.31) | **0.17 (2.35)** |
| Pattern comparison processing speed accuracy (NIH) | -0.01 (-0.24) | 0.05 (0.71) | **-0.14 (-2.19)** | -0.05 (-0.57) | -0.06 (-0.68) |
| Penn Progressive matrices accuracy | 0.01 (0.23) | **-0.20 (-2.58)** | -0.12 (-1.21) | **-0.17 (-2.02)** | 0.07 (0.74) |
| Penn Progressive matrices RT | 0.01 (0.09) | **-0.21 (-2.45)** | **-0.18 (-1.91)** | -0.12 (-1.28) | 0.07 (0.69) |
| Variable short Penn line orientation accuracy | 0.00 (-0.01) | -0.06 (-0.90) | -0.07 (-0.92) | **-0.14 (-1.78)** | 0.11 (1.30) |
| Variable short Penn line orientation RT | -0.05 (-0.77) | -0.05 (-0.67) | **-0.17 (-2.43)** | -0.06 (-0.65) | 0.02 (0.19) |
| Short Penn continuous performance true positives | **-0.09 (-1.83)** | -0.06 (-0.95) | 0.01 (0.16) | -0.05 (-0.71) | 0.04 (0.51) |
| Short Penn continuous performance sensitivity | -0.05 (-0.97) | 0.01 (0.14) | 0.07 (1.09) | -0.06 (-0.69) | 0.15 (2.02) |
| Short Penn continuous performance specificity | 0.04 (0.64) | -0.11 (-1.65) | -0.08 (-0.97) | -0.08 (-0.90) | 0.08 (0.89) |
| Penn word memory test accuracy | -0.05 (-1.03) | 0.07 (1.22) | -0.03 (-0.46) | -0.06 (-0.89) | -0.01 (-0.12) |
| Penn word memory test RT | -0.07 (-1.09) | 0.00 (-0.01) | 0.10 (1.31) | -0.10 (-1.17) | 0.03 (0.39) |
| List sorting working memory accuracy | -0.03 (-0.54) | **0.16 (2.02)** | -0.14 (-1.53) | **-0.16 (-1.82)** | 0.10 (1.02) |
| Language task accuracy | 0.00 (-0.01) | -0.02 (-0.20) | -0.06 (-0.69) | **-0.19 (-2.27)** | **0.17 (1.90)** |
| Language task RT | **-0.12 (-1.84)** | -0.02 (-0.22) | 0.03 (0.36) | **0.15 (1.83)** | -0.11 (-1.28) |
| Relational task accuracy | 0.05 (0.70) | -0.05 (-0.54) | -0.01 (-0.10) | -0.16 (-1.62) | **0.25 (2.90)** |
| Relational task RT | 0.06 (0.91) | -0.06 (-0.91) | -0.05 (-0.63) | 0.00 (0.04) | 0.01 (0.10) |
| Social task random stimuli rated random accuracy | 0.02 (0.34) | **-0.13 (-1.84)** | 0.05 (0.62) | **0.14 (1.75)** | -0.04 (-0.49) |
| Social task random stimuli rated random RT | 0.01 (0.25) | 0.12 (1.46) | -0.06 (-0.63) | **0.22 (2.52)** | 0.04 (0.43) |
| Social task social stimuli rated social accuracy | **0.12 (2.11)** | 0.01 (0.16) | -0.02 (-0.19) | **0.16 (2.27)** | 0.07 (0.86) |
| Social task social stimuli rated social RT | 0.01 (0.15) | **-0.13 (-1.71)** | -0.06 (-0.75) | **0.15 (1.79)** | 0.01 (0.09) |
| Working memory task accuracy | 0.10 (1.33) | -0.06 (-0.83) | -0.08 (-0.98) | -0.12 (-1.34) | **0.19 (2.39)** |
| Working memory task RT | **-0.12 (-2.06)** | 0.04 (0.54) | 0.05 (0.60) | 0.01 (0.15) | -0.12 (-1.26) |
| Delay discounting task $200 AUC | 0.13 (1.31) | **-0.18 (-2.30)** | -0.03 (-0.28) | **-0.22 (-2.18)** | 0.03 (0.30) |
| Delay discounting task $40K AUC | 0.07 (0.75) | **-0.16 (-1.90)** | -0.06 (-0.62) | -0.19 (-1.83) | 0.09 (0.82) |
| Gambling task larger prediction accuracy | -0.07 (-1.24) | -0.08 (-1.30) | 0.03 (0.37) | 0.04 (0.51) | -0.06 (-0.84) |
| Gambling task smaller prediction accuracy | 0.07 (1.24) | 0.08 (1.30) | -0.03 (-0.37) | -0.04 (-0.51) | 0.06 (0.84) |

| | | | | | |
|---|---|---|---|---|---|
| Gambling task larger prediction RT | -0.04 (-0.66) | -0.08 (-1.01) | 0.13 (1.60) | 0.04 (0.40) | -0.06 (-0.59) |
| Gambling task smaller prediction RT | -0.04 (-0.81) | -0.10 (-1.22) | 0.11 (1.27) | 0.07 (0.66) | 0.00 (0.04) |
| Penn emotion recognition Anger accuracy | -0.06 (-1.01) | 0.03 (0.56) | -0.02 (-0.26) | 0.06 (0.87) | 0.00 (-0.04) |
| Penn emotion recognition Fear accuracy | **-0.09 (-1.71)** | **0.14 (2.03)** | -0.08 (-1.05) | 0.09 (1.15) | 0.03 (0.32) |
| Penn emotion recognition Happy accuracy | **0.14 (1.80)** | 0.08 (1.06) | -0.12 (-1.45) | 0.04 (0.46) | 0.08 (0.96) |
| Penn emotion recognition Neutral accuracy | 0.02 (0.45) | 0.03 (0.44) | **0.10 (1.74)** | 0.04 (0.57) | 0.04 (0.59) |
| Penn emotion recognition Sad accuracy | -0.04 (-0.71) | 0.03 (0.34) | **-0.17 (-2.75)** | -0.01 (-0.14) | -0.01 (-0.10) |
| Penn emotion recognition Total accuracy | -0.06 (-1.11) | 0.11 (1.65) | -0.08 (-1.13) | 0.08 (1.16) | 0.04 (0.48) |
| Penn emotion recognition Total RT | -0.03 (-0.46) | 0.00 (0.00) | -0.09 (-1.07) | -0.06 (-0.57) | 0.11 (1.05) |
| Hariri Emotion task accuracy | 0.03 (0.44) | 0.02 (0.28) | -0.08 (-0.96) | **-0.17 (-2.30)** | -0.01 (-0.13) |
| Hariri Emotion task RT | -0.03 (-0.50) | -0.05 (-0.66) | 0.00 (-0.04) | **0.19 (1.90)** | 0.00 (-0.05) |
| Anger affect (NIH) | **-0.44 (-3.68)** | **-0.33 (-4.08)** | -0.08 (-0.66) | 0.00 (0.01) | 0.02 (0.26) |
| Anger hostility (NIH) | **-0.32 (-3.19)** | **-0.17 (-2.28)** | -0.07 (-0.77) | 0.01 (0.14) | -0.12 (-1.57) |
| Anger physical aggression (NIH) | **-0.25 (-3.71)** | -0.06 (-0.83) | 0.02 (0.29) | 0.05 (0.63) | 0.03 (0.40) |
| Fear affect (NIH) | **-0.51 (-3.89)** | **-0.33 (-3.93)** | 0.03 (0.27) | -0.08 (-0.74) | -0.12 (-1.24) |
| Fear somatic arousal (NIH) | **-0.44 (-3.75)** | **-0.22 (-2.50)** | -0.08 (-0.76) | 0.14 (1.41) | -0.11 (-1.18) |
| Sadness (NIH) | **-0.40 (-3.50)** | **-0.32 (-3.73)** | -0.12 (-0.97) | 0.01 (0.11) | -0.11 (-1.18) |
| Life satisfaction (NIH) | **0.35 (3.38)** | **0.20 (2.58)** | 0.10 (1.04) | -0.08 (-0.95) | -0.01 (-0.16) |
| Meaning purpose (NIH) | **0.37 (4.62)** | 0.09 (1.14) | 0.10 (1.23) | 0.06 (0.71) | -0.10 (-1.22) |
| Positive affect (NIH) | **0.34 (3.97)** | **0.20 (2.39)** | 0.13 (1.39) | -0.07 (-0.74) | -0.10 (-1.17) |
| Friendship (NIH) | **0.20 (2.77)** | **0.23 (2.40)** | **0.22 (2.14)** | 0.11 (1.14) | 0.08 (0.87) |
| Loneliness (NIH) | **-0.34 (-3.71)** | **-0.22 (-2.69)** | -0.13 (-1.30) | -0.01 (-0.05) | -0.08 (-0.90) |
| Perceived hostility (NIH) | **-0.22 (-2.78)** | **-0.18 (-2.07)** | -0.10 (-1.06) | -0.03 (-0.25) | **-0.17 (-1.71)** |
| Perceived rejection (NIH) | **-0.31 (-3.75)** | **-0.17 (-1.77)** | **-0.20 (-2.19)** | 0.02 (0.21) | -0.12 (-1.29) |
| Emotional support (NIH) | **0.25 (3.69)** | 0.05 (0.54) | **0.18 (2.68)** | 0.06 (0.72) | -0.02 (-0.18) |
| Instrumental support (NIH) | **0.16 (2.67)** | 0.08 (1.20) | 0.11 (1.60) | 0.02 (0.24) | 0.02 (0.27) |
| Perceived stress (NIH) | **-0.46 (-3.38)** | **-0.36 (-4.15)** | -0.07 (-0.53) | 0.05 (0.39) | -0.09 (-0.90) |
| Self-efficacy (NIH) | **0.17 (2.80)** | **0.17 (2.05)** | 0.06 (0.60) | 0.15 (1.60) | 0.08 (0.82) |
| Agreeableness (NEO-FFI) | **0.21 (2.22)** | 0.00 (-0.03) | 0.05 (0.52) | **-0.22 (-2.62)** | -0.01 (-0.13) |
| Openness experience (NEO-FFI) | **-0.13 (-2.08)** | -0.10 (-1.13) | **-0.22 (-2.94)** | 0.03 (0.28) | -0.08 (-0.94) |
| Conscientiousness (NEO-FFI) | **0.28 (4.00)** | **0.34 (3.45)** | 0.18 (1.41) | 0.02 (0.20) | **-0.23 (-2.40)** |
| Neuroticism (NEO-FFI) | **-0.48 (-3.79)** | **-0.27 (-3.26)** | -0.05 (-0.47) | 0.08 (0.79) | -0.08 (-0.92) |
| Extraversion (NEO-FFI) | **0.20 (2.97)** | **0.13 (1.74)** | **0.13 (1.72)** | 0.08 (0.95) | -0.03 (-0.32) |
| Pain interference (NIH) | **-0.37 (-3.63)** | -0.09 (-1.08) | 0.08 (0.86) | -0.07 (-0.70) | **-0.16 (-1.87)** |
| Lifetime alcohol dependence symptoms (SSAGA) | **-0.24 (-3.27)** | -0.09 (-1.02) | 0.00 (0.03) | **-0.20 (-2.28)** | **-0.19 (-2.23)** |
| Lifetime alcohol abuse diagnosis (SSAGA) | **-0.18 (-2.08)** | **-0.14 (-1.93)** | 0.04 (0.47) | 0.04 (0.44) | -0.08 (-0.92) |
| Lifetime alcohol abuse symptoms (SSAGA) | **-0.25 (-3.11)** | -0.12 (-1.46) | 0.06 (0.64) | -0.10 (-1.20) | -0.09 (-1.02) |
| Lifetime alcohol dependence diagnosis (SSAGA) | **-0.15 (-2.45)** | -0.09 (-1.09) | 0.00 (-0.04) | **-0.26 (-3.16)** | **-0.17 (-1.89)** |
| Drinks per day past 12 months (SSAGA) | **-0.18 (-3.02)** | 0.08 (1.16) | 0.02 (0.22) | -0.03 (-0.39) | 0.06 (0.81) |
| Frequency alcohol use past 12 months (SSAGA) | **0.10 (1.86)** | -0.02 (-0.23) | -0.05 (-0.75) | 0.05 (0.60) | 0.12 (1.61) |
| Frequency drunk past 12 months (SSAGA) | **0.16 (2.93)** | 0.10 (1.24) | -0.13 (-1.62) | **0.14 (1.69)** | 0.02 (0.29) |
| Age first alcohol use (SSAGA) | **0.22 (3.32)** | 0.02 (0.27) | -0.07 (-0.98) | 0.11 (1.50) | -0.05 (-0.58) |
| Drinks per day heaviest period (SSAGA) | **-0.14 (-2.20)** | 0.05 (0.70) | 0.11 (1.64) | -0.01 (-0.10) | 0.02 (0.30) |
| Frequency alcohol use heaviest period (SSAGA) | **0.10 (1.77)** | 0.02 (0.37) | -0.01 (-0.15) | 0.07 (1.04) | 0.07 (0.99) |
| Frequency drunk heaviest period (SSAGA) | **0.12 (1.69)** | 0.07 (0.96) | -0.06 (-0.83) | 0.03 (0.31) | 0.02 (0.23) |
| History of smoking (SSAGA) | **-0.15 (-2.18)** | -0.05 (-0.76) | 0.02 (0.34) | 0.01 (0.07) | -0.06 (-0.79) |
| Currently smoking (SSAGA) | **-0.16 (-1.99)** | -0.01 (-0.16) | 0.13 (1.46) | 0.13 (1.39) | **-0.19 (-2.26)** |
| Times used nonmarijuana illicit drugs (SSAGA) | **-0.21 (-3.05)** | **-0.20 (-2.60)** | 0.11 (1.19) | -0.04 (-0.44) | 0.03 (0.38) |
| Times used cocaine (SSAGA) | **-0.21 (-2.87)** | -0.10 (-1.50) | 0.03 (0.39) | -0.01 (-0.08) | 0.00 (0.02) |
| Times used hallucinogens (SSAGA) | **-0.11 (-1.77)** | **-0.21 (-3.00)** | 0.06 (0.61) | -0.04 (-0.45) | -0.03 (-0.39) |

| | | | | | |
|---|---|---|---|---|---|
| Times used opiates (SSAGA) | **-0.19 (-2.73)** | **-0.20 (-2.53)** | 0.12 (1.14) | 0.01 (0.14) | -0.08 (-0.90) |
| Times used sedatives (SSAGA) | **-0.27 (-3.37)** | **-0.13 (-1.82)** | 0.02 (0.18) | 0.02 (0.28) | -0.02 (-0.30) |
| Times used stimulants (SSAGA) | -0.10 (-1.64) | **-0.12 (-1.69)** | 0.01 (0.19) | -0.04 (-0.49) | 0.03 (0.41) |
| Times used marijuana (SSAGA) | **-0.17 (-2.56)** | -0.08 (-1.15) | 0.08 (1.09) | -0.07 (-0.92) | -0.05 (-0.67) |
| Marijuana dependence (SSAGA) | -0.09 (-1.38) | **-0.28 (-3.92)** | 0.06 (0.60) | -0.04 (-0.47) | 0.05 (0.59) |

**Table S3**. Cross-validated canonical correlation analysis. Significant p-values are indicated in bold.

| | In-sample correlation | | Out-of-sample correlation | | |
|---|---|---|---|---|---|
| | Whole set | Permuted p | Training sets (mean [range] across folds) | Test sets (mean [range] across folds) | Permuted p (mean [range] across folds) |
| LC1 | *r* = 0.69 | ***p* = 0.0002** | r=0.71 [0.71-0.72] | *r* = 0.49 [0.44-0.53] | ***p* = 0.001 [0.001-0.001]** |
| LC2 | *r* = 0.53 | ***p* = 0.0002** | r=0.57 [0.55-0.58] | *r* = 0.19 [0.11-0.26] | ***p* = 0.039 [0.001-0.155]** |
| LC3 | *r* = 0.49 | ***p* = 0.0002** | r=0.54 [0.53-0.55] | *r* = 0.12 [0.06-0.29] | *p* = 0.209 [0.001-0.398] |
| LC4 | *r* = 0.44 | ***p* = 0.0002** | r=0.49 [0.47-0.51] | *r* = 0.01 [-0.05-0.07] | *p* = 0.497 [0.169-0.873] |
| LC5 | *r* = 0.42 | ***p* = 0.0002** | r=0.46 [0.44-0.48] | *r* = 0.06 [-0.09-0.24] | *p* = 0.336 [0.017-0.787] |

**Table S4**. Post-hoc associations between sleep (or biopsychosocial; BPS) composite scores and sociodemographic measures, as well as physical and mental health measures. Associations using continuous measures (e.g., age, education years) were tested with Pearson correlations, while categorical measures (e.g., sex, race) were assessed with t-tests or analyses of variance. Associations indicated in bold were found to be statistically significant after FDR correction ($q < 0.05$). Tests with <5 subjects in a category were not computed because of too little variance.

| | | LC1 | | LC2 | | LC3 | | LC4 | | LC5 | |
|---|---|---|---|---|---|---|---|---|---|---|---|
| | | Sleep scores | BPS scores | Sleep scores | BPS scores | Sleep scores | BPS scores | Sleep scores | BPS scores | Sleep scores | BPS scores |
| Age | *r* | 0.02 | 0.01 | -0.01 | -0.02 | -0.04 | -0.06 | **0.08** | **0.10** | 0.03 | -0.03 |
| | *p* | 0.492 | 0.851 | 0.790 | 0.621 | 0.308 | 0.085 | **0.020** | **0.007** | 0.398 | 0.478 |
| Sex | *T* | -0.86 | -0.86 | 0.52 | 0.51 | -0.92 | -0.92 | 1.75 | 1.88 | **-2.88** | **-4.35** |
| | *p* | 0.388 | 0.388 | 0.601 | 0.613 | 0.359 | 0.357 | 0.081 | 0.061 | **0.004** | **0.000** |
| Education | *r* | **-0.11** | **-0.14** | **0.10** | **0.11** | 0.00 | -0.03 | **-0.07** | **-0.12** | **-0.11** | **-0.22** |
| | *p* | **0.003** | **0.000** | **0.007** | **0.002** | 0.918 | 0.379 | **0.048** | **0.001** | **0.002** | **0.000** |
| Race | *F* | 0.69 | 1.07 | 0.24 | 1.32 | **3.76** | 1.19 | **2.44** | **3.22** | **2.67** | **4.79** |
| | *p* | 0.631 | 0.373 | 0.945 | 0.254 | **0.002** | 0.312 | **0.033** | **0.007** | **0.021** | **0.000** |
| Ethnicity | *F* | 2.11 | **5.29** | 0.25 | 1.79 | 0.83 | 1.57 | 0.66 | 1.72 | 0.50 | 1.36 |
| | *p* | 0.122 | **0.005** | 0.780 | 0.168 | 0.436 | 0.209 | 0.517 | 0.180 | 0.608 | 0.257 |
| Household income | *F* | **2.35** | **3.82** | 0.89 | **2.69** | 0.52 | 0.55 | 0.83 | 0.89 | 1.80 | 0.88 |
| | *p* | **0.022** | **0.000** | 0.513 | **0.009** | 0.817 | 0.798 | 0.565 | 0.512 | 0.083 | 0.518 |
| Employment status | *F* | 1.89 | 2.42 | **3.95** | 1.34 | 0.23 | 1.08 | 2.96 | **4.41** | 0.16 | 0.17 |
| | *p* | 0.152 | 0.090 | **0.020** | 0.262 | 0.796 | 0.342 | 0.052 | **0.013** | 0.849 | 0.840 |
| School status | *F* | -0.10 | -0.18 | **-2.23** | -1.46 | -0.80 | 1.18 | **-2.11** | -0.86 | -0.42 | -1.19 |
| | *p* | 0.923 | 0.857 | **0.026** | 0.146 | 0.424 | 0.237 | **0.035** | 0.388 | 0.673 | 0.236 |
| Relationship status | *F* | 0.43 | 1.05 | -0.06 | 0.23 | 1.16 | 0.64 | -0.15 | 0.78 | -0.68 | 0.40 |
| | *p* | 0.665 | 0.294 | 0.955 | 0.816 | 0.247 | 0.523 | 0.882 | 0.437 | 0.495 | 0.689 |
| BMI | *r* | 0.04 | 0.03 | -0.05 | -0.02 | **-0.11** | -0.03 | **0.08** | 0.05 | 0.05 | -0.01 |
| | *p* | 0.261 | 0.463 | 0.151 | 0.603 | **0.003** | 0.394 | **0.024** | 0.196 | 0.179 | 0.678 |
| Hematocrit | *r* | 0.05 | 0.02 | -0.03 | 0.01 | -0.03 | -0.04 | -0.02 | 0.01 | -0.02 | 0.05 |
| | *p* | 0.150 | 0.499 | 0.423 | 0.819 | 0.461 | 0.214 | 0.514 | 0.831 | 0.652 | 0.142 |
| Blood pressure | *F* | 0.87 | 0.57 | 1.09 | 0.46 | 0.86 | 0.53 | 1.57 | 0.46 | 0.82 | 0.48 |
| | *p* | 0.483 | 0.683 | 0.360 | 0.768 | 0.485 | 0.714 | 0.179 | 0.767 | 0.514 | 0.747 |
| Maternal history of depression | *t* | -1.50 | -1.76 | -1.54 | -1.39 | -0.36 | 0.78 | 1.22 | 0.27 | -0.20 | -1.75 |
| | *p* | 0.135 | 0.080 | 0.123 | 0.166 | 0.716 | 0.438 | 0.222 | 0.791 | 0.840 | 0.080 |
| Paternal history of depression | *t* | -1.44 | **-2.01** | **-2.24** | -0.80 | 0.54 | -1.00 | -0.76 | 1.48 | -1.18 | -1.65 |
| | *p* | 0.150 | **0.044** | **0.025** | 0.425 | 0.587 | 0.320 | 0.447 | 0.139 | 0.238 | 0.100 |
| Maternal history of BD | *t* | -0.18 | 0.79 | -1.37 | -0.64 | -0.17 | -1.03 | -1.84 | -0.35 | -1.17 | -1.31 |
| | *p* | 0.860 | 0.428 | 0.173 | 0.522 | 0.868 | 0.303 | 0.066 | 0.726 | 0.242 | 0.190 |
| Paternal history of BD | *t* | -0.39 | -0.23 | -1.21 | -0.54 | 1.93 | 1.69 | -1.05 | 1.85 | 0.42 | -0.76 |
| | *p* | 0.694 | 0.815 | 0.228 | 0.590 | 0.055 | 0.091 | 0.293 | 0.065 | 0.677 | 0.450 |
| Maternal history of anxiety | *t* | -1.82 | -0.86 | -1.38 | **-2.38** | -0.25 | -1.25 | 0.62 | 0.34 | 0.05 | -1.79 |
| | *p* | 0.069 | 0.391 | 0.167 | **0.017** | 0.801 | 0.213 | 0.532 | 0.735 | 0.959 | 0.074 |
| Paternal history of anxiety | *t* | -0.67 | 0.38 | -1.54 | -0.52 | 1.13 | 0.39 | 0.85 | **2.22** | -0.92 | 0.07 |
| | *p* | 0.505 | 0.708 | 0.124 | 0.606 | 0.261 | 0.693 | 0.397 | **0.027** | 0.357 | 0.948 |
| Maternal history of drug or | *t* | -0.91 | -0.76 | 0.91 | 0.59 | -0.22 | 0.92 | 0.02 | -1.97 | -1.47 | -1.70 |
| | *p* | 0.366 | 0.448 | 0.363 | 0.556 | 0.829 | 0.357 | 0.987 | 0.050 | 0.143 | 0.089 |

| | | | | | | | | | | | |
|---|---|---|---|---|---|---|---|---|---|---|---|
| alcohol problems | | | | | | | | | | | |
| Paternal history of drug or alcohol problems | *t* | -0.61 | -1.05 | -0.90 | **-2.21** | 1.08 | 1.57 | -0.60 | -0.16 | -0.11 | -0.95 |
| | *p* | 0.542 | 0.293 | 0.369 | **0.028** | 0.283 | 0.117 | 0.549 | 0.870 | 0.916 | 0.343 |
| Maternal history of SZ | *t* | - | - | - | - | - | - | - | - | - | - |
| | *p* | - | - | - | - | - | - | - | - | - | - |
| Paternal history of SZ | *t* | - | - | - | - | - | - | - | - | - | - |
| | *p* | - | - | - | - | - | - | - | - | - | - |
| Maternal history of Alzheimer's disease or dementia | *t* | - | - | - | - | - | - | - | - | - | - |
| | *p* | - | - | - | - | - | - | - | - | - | - |
| Paternal history of Alzheimer's disease or dementia | *t* | - | - | - | - | - | - | - | - | - | - |
| | *p* | - | - | - | - | - | - | - | - | - | - |
| Maternal history of Parkinson's disease | *t* | - | - | - | - | - | - | - | - | - | - |
| | *p* | - | - | - | - | - | - | - | - | - | - |
| Paternal history of Parkinson's disease | *t* | - | - | - | - | - | - | - | - | - | - |
| | *p* | - | - | - | - | - | - | - | - | - | - |
| Maternal history of Tourette's syndrome | *t* | - | - | - | - | - | - | - | - | - | - |
| | *p* | - | - | - | - | - | - | - | - | - | - |
| Paternal history of Tourette's syndrome | *t* | - | - | - | - | - | - | - | - | - | - |
| | *p* | - | - | - | - | - | - | - | - | - | - |

**Table S5**. Correlations between sleep (or biopsychosocial) loadings in control analyses with those from the original analysis.

| | | Not regressing out confounds | Excluding participants positive for substance use | Excluding physical health variables * | Excluding demographic variables** | Quantile normalization on biopsychosocial / sleep data |
|---|---|---|---|---|---|---|
| LC1 | Sleep loadings | -1.00 | -0.97 | -1.00 | -1.00 | 0.62 |
| | Biopsychosocial loadings | -1.00 | -0.99 | -1.00 | -1.00 | -0.16 |
| LC2 | Sleep loadings | -1.00 | -0.96 | -0.99 | -1.00 | 0.95 |
| | Biopsychosocial loadings | -0.99 | -0.95 | -1.00 | -1.00 | 0.97 |
| LC3 | Sleep loadings | 1.00 | 0.93 | 0.99 | 1.00 | -0.71 |
| | Biopsychosocial loadings | 0.99 | 0.85 | 0.97 | 0.99 | -0.58 |
| LC4 | Sleep loadings | 0.98 | 0.81 | 0.98 | 0.98 | -0.95 |
| | Biopsychosocial loadings | 0.94 | 0.75 | 0.99 | 0.97 | -0.89 |
| LC5 | Sleep loadings | -0.94 | -0.59 | -0.87 | 0.95 | 0.53 |
| | Biopsychosocial loadings | -0.94 | -0.58 | -0.91 | 0.95 | 0.67 |

* BMI, Hematocrit, Blood pressure

** Employment status, Household income, In school, Relationship status

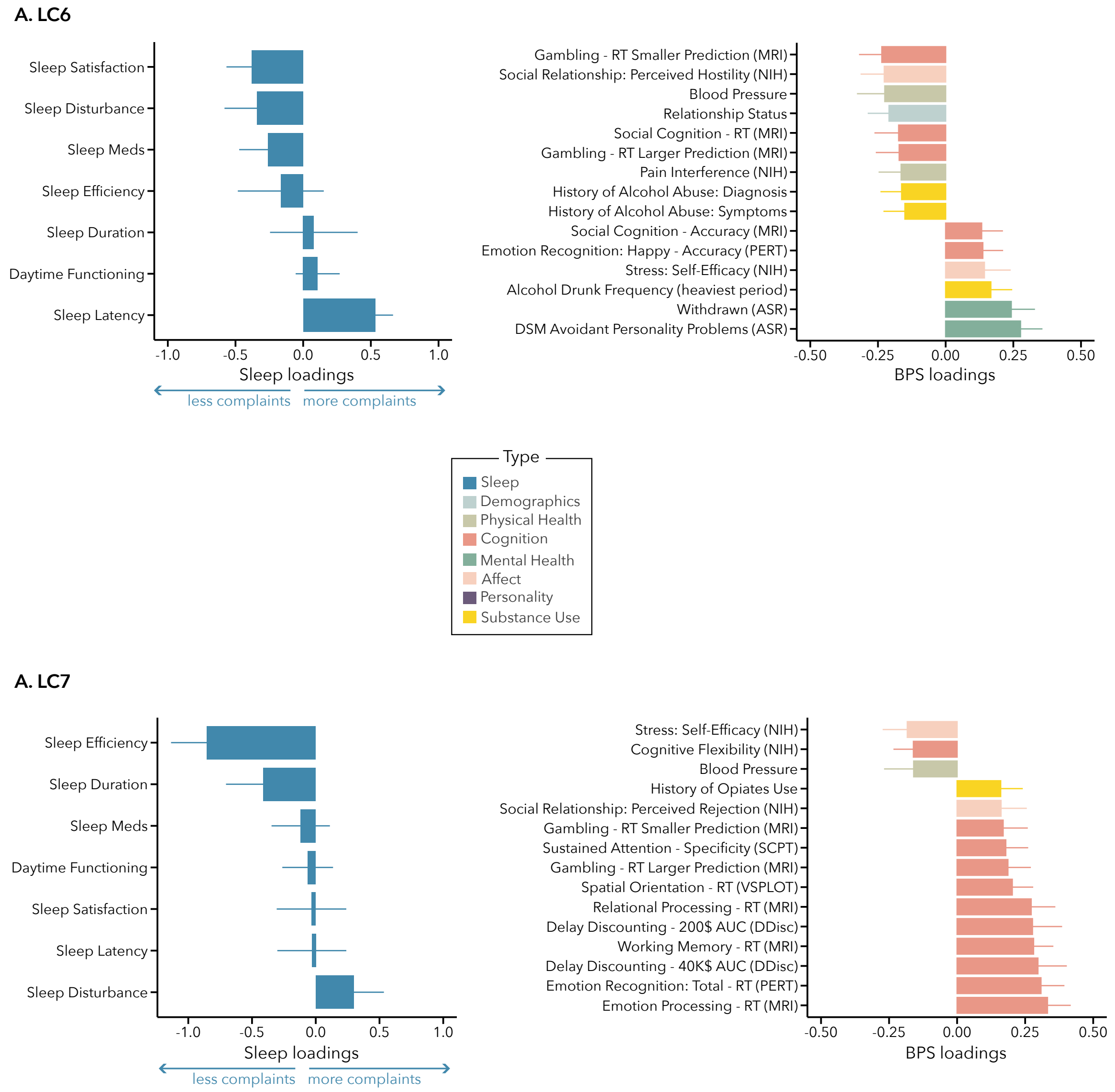


**Figure S1**. Latent Components LC6 and LC7

*(A) Sleep loadings (left) and top 15 strongest biopsychosocial (BPS) loadings (right) for LC6. (B) Sleep loadings (left) and top 15 strongest biopsychosocial (BPS) loadings (right) for LC7.*

*Positive values on sleep (blue) loadings indicate worse outcomes while positive values on biopsychosocial loadings reflect higher magnitude on these measures. Error bars indicate bootstrapped-estimated confidence intervals (i.e., standard deviation) and measures in bold indicate statistical significance*

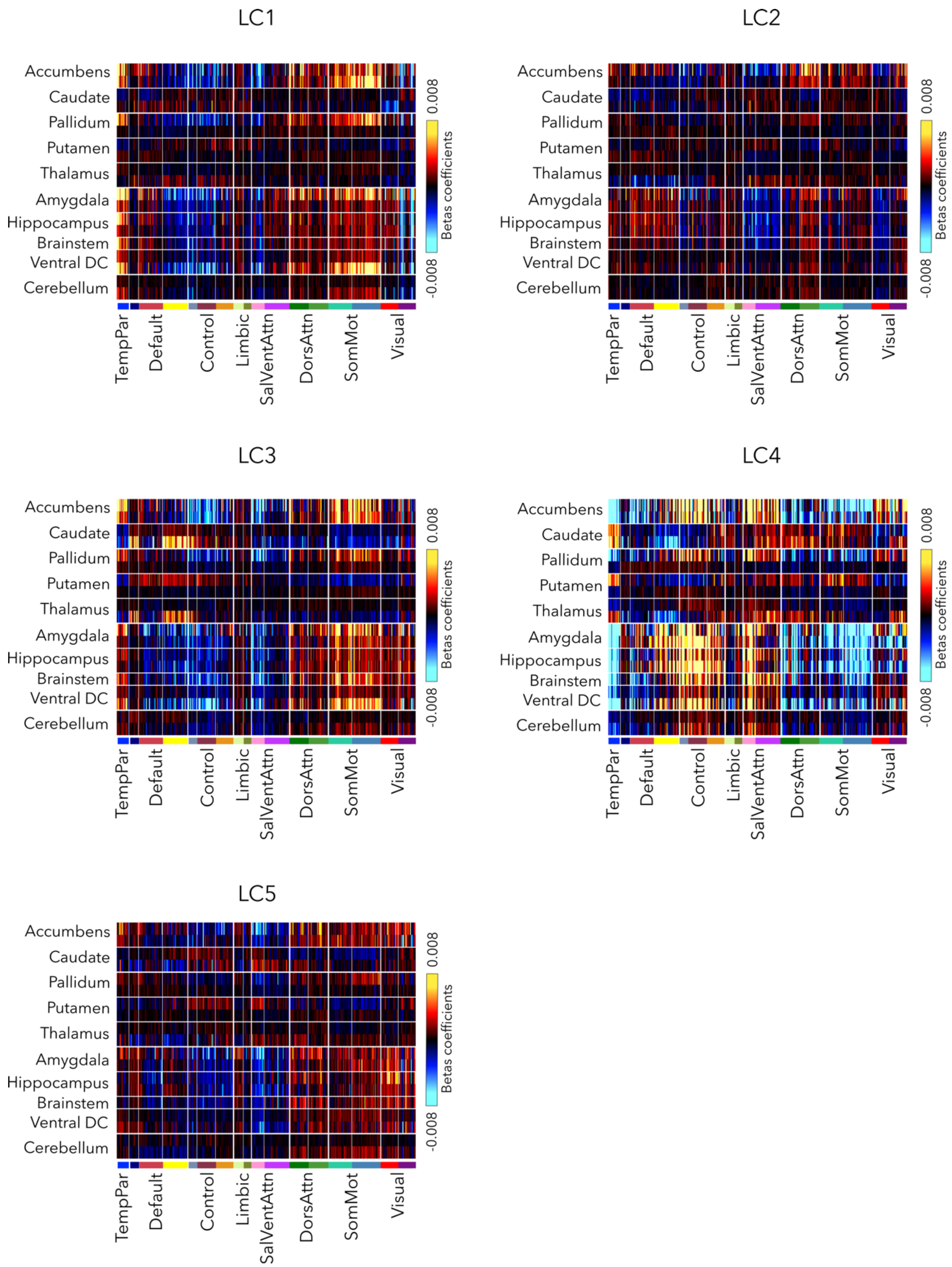


**Figure S2**. Post-hoc associations between CCA mean composite scores and RSFC, highlighting beta coefficients between subcortical regions and cortical networks.

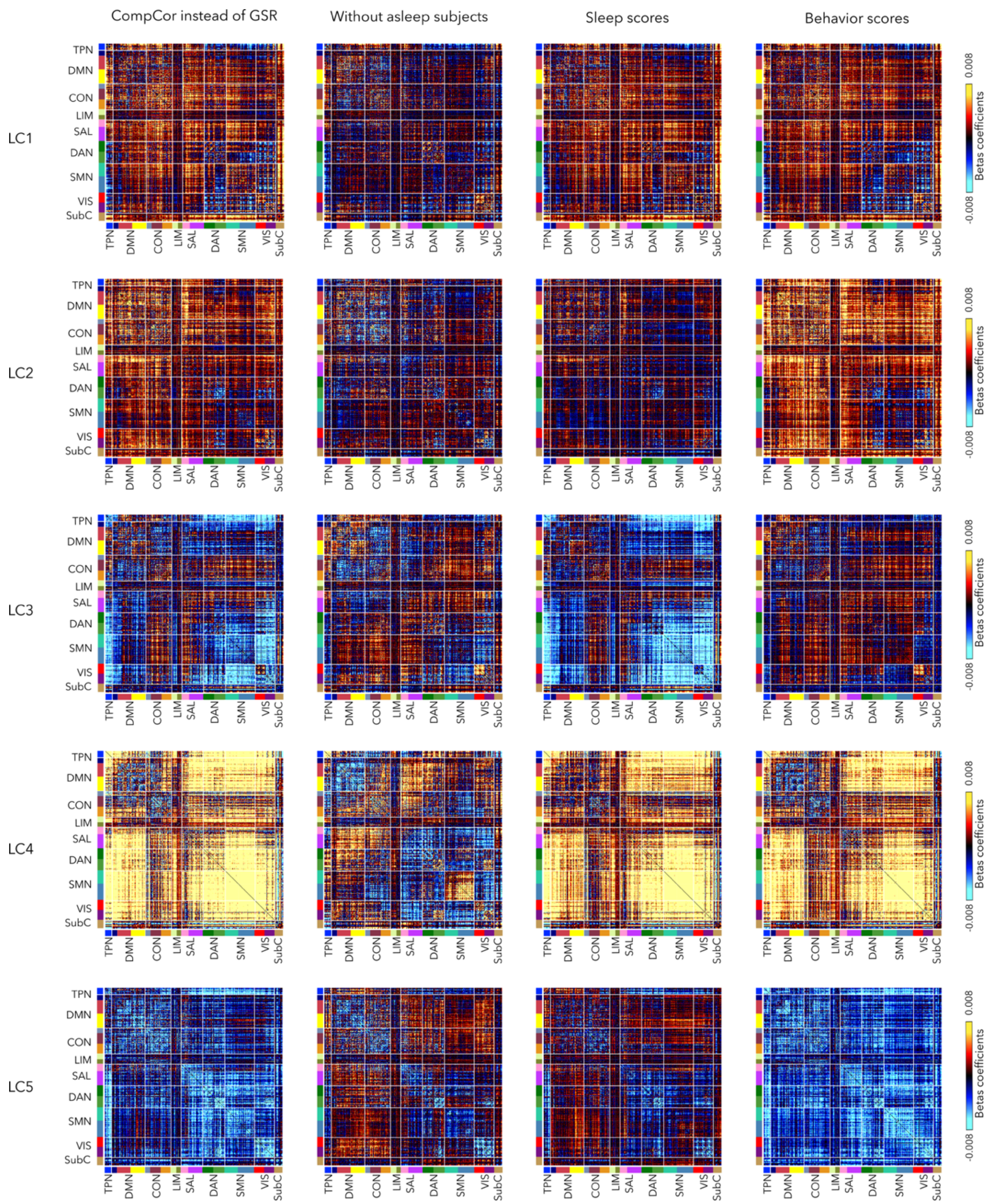


**Figure S3**. Control analyses for post-hoc associations with RSFC. (**Left panel**) GLM analysis using RSFC data that underwent CompCor instead of global signal regression. (**Middle left panel**) GLM analysis after excluding subjects that likely fell asleep in the scanner (N=100). (**Middle right panel**) GLM analysis between RSFC and *sleep* composite scores (instead of averaged composite scores). (**Right panel**) GLM analysis between RSFC and *biopsychosocial* composite scores (instead of averaged composite scores).

*Abbreviations: CON=Executive control network, DAN=Dorsal attention network, DMN=Default mode network, LIM=Limbic network, SAL=Salience/Ventral attention network, SMN=Somatosensory-motor network, SubC=Subcortical regions, TPN=Temporoparietal network, VIS=Visual network.*

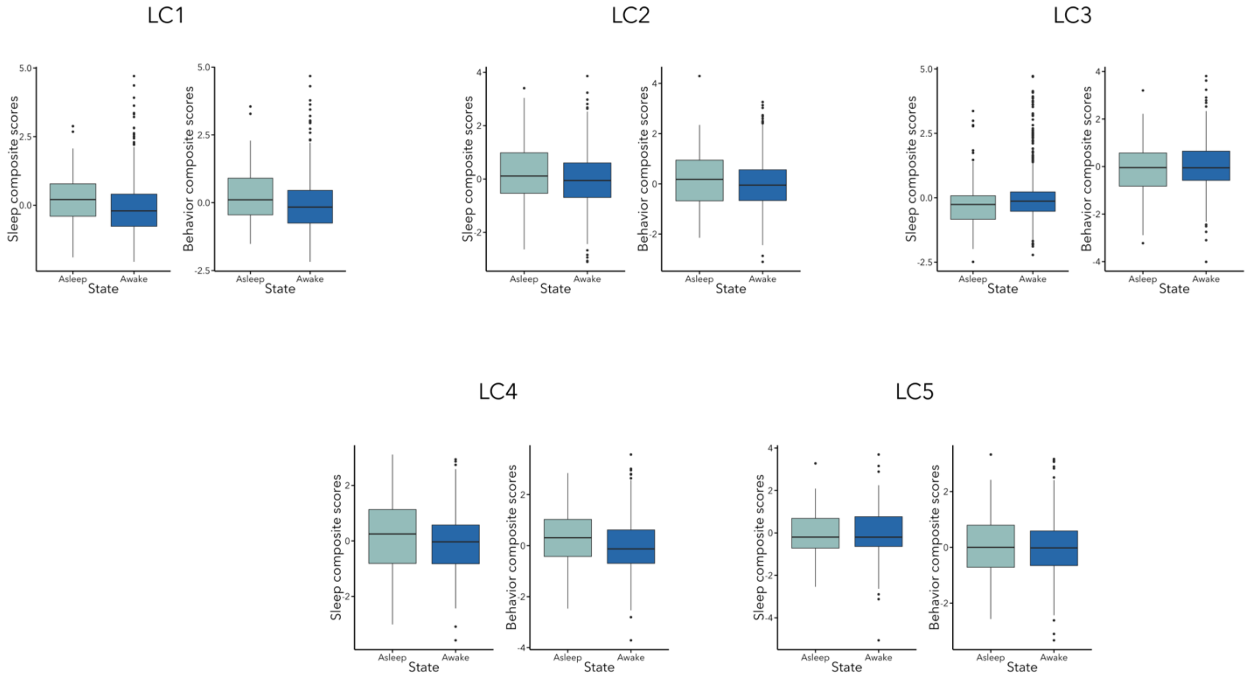


**Figure S4**. Post-hoc t-tests assessing differences in sleep (or biopsychosocial) composite scores between participants who likely stayed awake (N=623) in the scanner vs. those who likely fell asleep in the scanner (N=100).

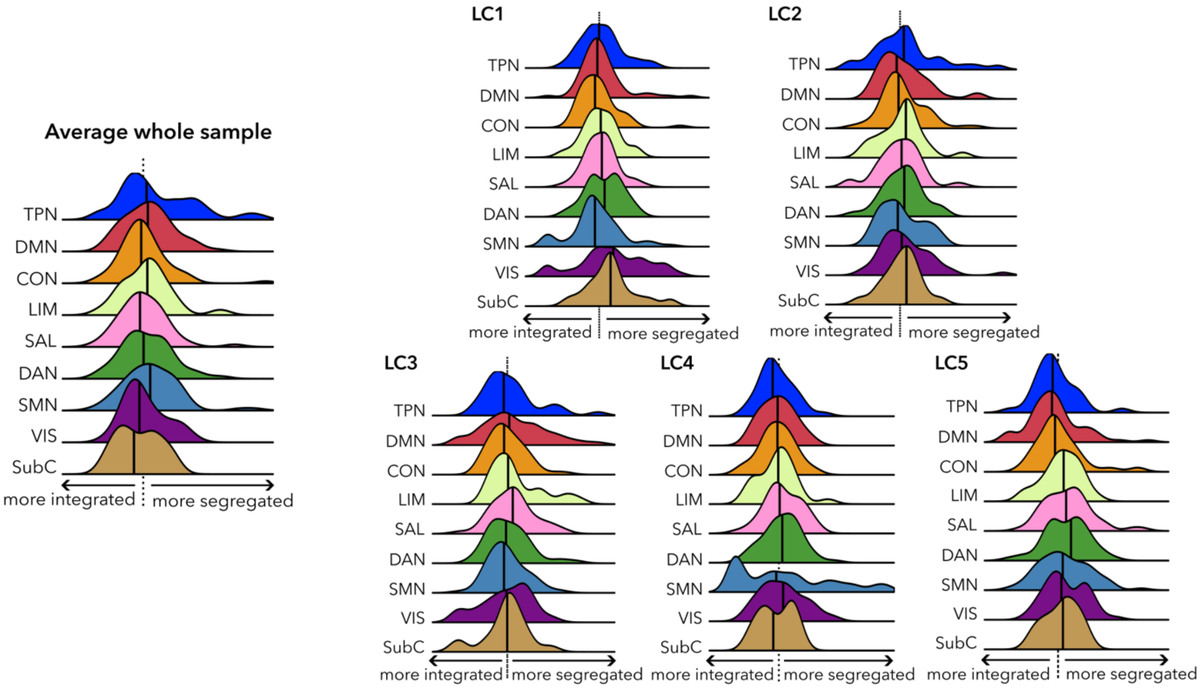


**Figure S5**. Distribution of the segregation and integration ratio on the average RSFC matrix of the whole sample compared to integration and segregation measures per LC. The dashed line indicates the median of all parcels, and the bold black lines represent the median for each network.

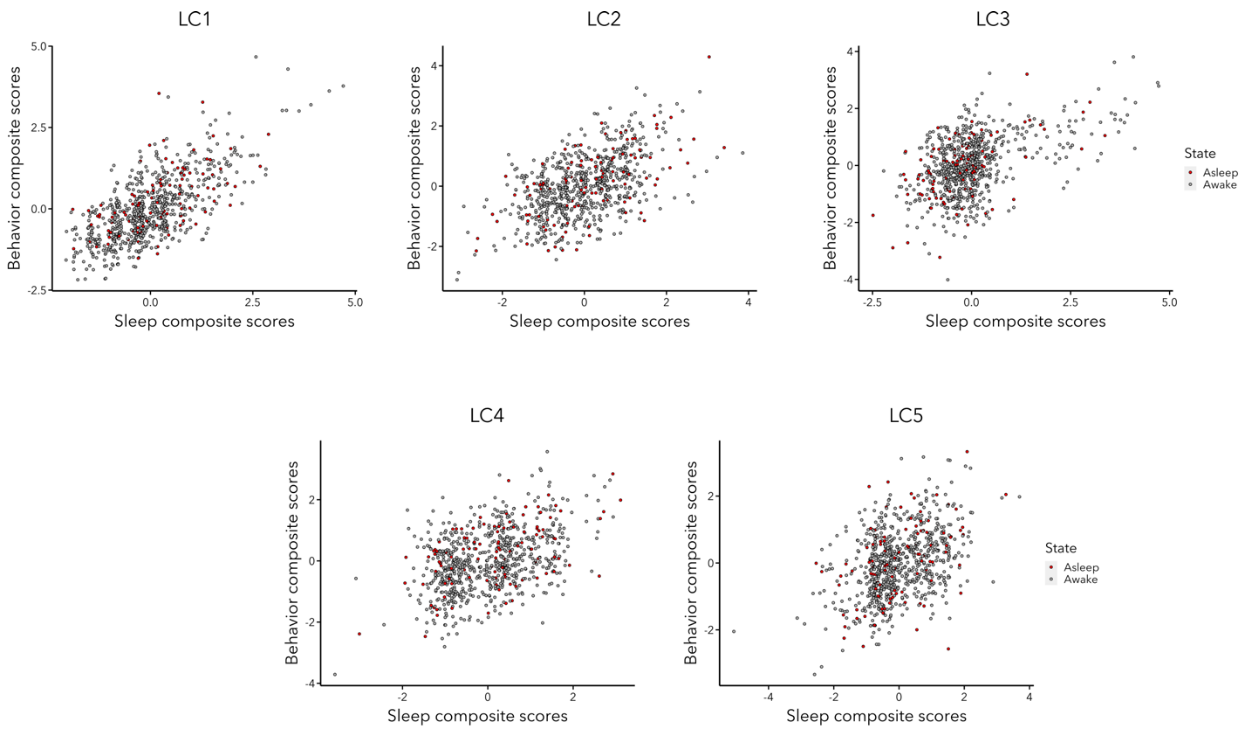


**Figure S6**. Scatterplots showing the distribution of sleep and biopsychosocial composite scores of participants who likely fell asleep in the scanner (in red) compared to those that likely stayed awake (in grey).
